# Complex life cycles and parasite-mediated trophic cascades drive species coexistence and the maintenance of genetic diversity

**DOI:** 10.1101/2023.12.22.571902

**Authors:** Ana C. Hijar-Islas, Nathaniel V. Mon Père, Weini Huang, Christophe Eizaguirre

## Abstract

Ecological and evolutionary processes shape the dynamics of species interactions, yet the drivers of eco-evolutionary feedback remain elusive. Here, we developed an individual-based model of a coevolving predator-prey-parasite system, in which predators can be infected by trophically transmitted parasites. We combined host-parasite coevolution with prey-predator interactions at an individual level, hence integrating evolutionary and ecological processes. We show that species coexist more when parasite-mediated selection is weak on both the predator and the prey. Population size and genotypic diversity of the parasite are highest at intermediate parasite-mediated selection on the predator and no parasite-mediated selection on the prey. Interestingly, we show that the evolution of super-resistant genotypes, where host genotypes have resistant alleles in all loci, is driven by interspecific interactions rather than by the population size of the host species. Overall, host-parasite coevolution shapes the direction of interspecific ecological interactions which, in turn, determine species coexistence and community diversity in a complex system.

## Introduction

Parasites are arguably one of the strongest selection pressures, as they drive the evolution of key traits in their hosts, including genetic diversity (Schmid-Hempel, 2021). Parasites also play a key role in ecology and community dynamics being important consumers in food webs (Lafferty & Kuris, 2002). Their biomass can exceed that of top predators (Kuris et al., 2008), and most food web links involve parasite species (Lafferty et al., 2008). They are involved in food web stability and interactions via density-mediated trophic cascades (Brunner et al. 2017) and host manipulation that facilitate their transmission (Lion et al., 2006). In turn, some parasites depend on trophic links to complete their complex life cycles (Lafferty et al., 2008). While often compared, parasite pressure is different from predation, as a predator can consume multiple whole prey individuals and a parasite mostly feeds from a single host individual (Lafferty & Kuris, 2002).

Host-parasite coevolution can be driven by fluctuating selection dynamics, where multiple genotypes coexist, or by arms race dynamics, where recurrent novel genotypes reach near fixation in the population (Tellier et al., 2014). Under the arms race predictions, a genotype’s advantage is temporary and depends on the abundance and genotype of the antagonistic counterpart (Retel et al., 2019). From a theoretical point of view, host-parasite coevolution is often modelled using a gene-for-gene (GFG) framework (Thompson & Burdon, 1992; Sasaki, 2000; Segarra, 2005). In this framework, some parasite genotypes can infect a wider range of hosts than others, with certain genotypes exhibiting universal virulence (Buckingham & Ashby, 2022). On the other hand, some hosts are better equipped to fend off a greater variety of parasites (Buckingham & Ashby, 2022). The GFG framework enables the exploration of the emergence of host and parasite supertypes, in which fitness is maximised.

Recent theoretical work shows that population dynamics and specialism drive host-parasite coevolution (Buckingham & Ashby, 2022). Fluctuating selection generates host and parasite diversity, and it is typically attributed to highly specific host-parasite interactions (Agrawal & Lively, 2002). However, evidence also suggests that specificity from the matching-alleles framework is not necessarily required to generate fluctuating selection dynamics and that ecological dynamics are important (Best et al. 2017). Population dynamics can have significant effects on coevolutionary dynamics (Buckingham & Ashby, 2022) because population sizes ultimately determine the supply of *de novo* mutations (Retel et al., 2019). Particularly, in small population sizes, demographic fluctuations and genetic drift may lead to extinction (Huang et al., 2015) ending the coevolutionary process.

In natural systems, host-parasite coevolution does not occur in isolation, and both predator and prey can become hosts (Lafferty, 1992; Strona, 2015; Brunner et al., 2017). We are interested in how the entangled predator-prey and host-parasite interactions can impact the evolution of all species and their ecological dynamics. In predator-prey dynamics, experimental evidence shows that the rapid evolution of predators’ attack traits and prey’s defence traits can influence population dynamics (Yoshida et al., 2003; Hairston Jr. et al., 2005; Becks et al., 2012). Theory further predicts that when predators’ attack traits evolve faster than prey’s defence traits, predator-prey dynamics result in large-amplitude oscillations of population size (Mougi & Iwasa, 2010), which can increase the likelihood of extinction. Contrarily, when prey’s defence traits evolve faster than predators’ attack traits, either a stable equilibrium or small amplitude oscillations of population size are likely to occur. Opposite to predator-prey systems, parasites typically evolve faster than the host due to their short generation times (Mougi & Iwasa, 2010). When multiple parasite species attack a single host, host-parasite dynamics tend to have large-amplitude oscillations (Dawkins & Krebs, 1979; Mougi & Iwasa, 2010). In an experimental setup where predator-prey and host-parasite interactions are integrated, the evolution of prey resistance can maintain species coexistence by allowing the prey population to support both consumers (Frickel et al., 2017). Yet, the influence of multiple species evolution on species coexistence and the maintenance of genetic diversity remains to be fully explored in a general framework.

Individual-based modelling is a powerful tool to model such a complex system with integrated coevolving species and allow eco-evolutionary feedback because mutation and extinction happen in individual reproduction and death events. When species interactions and coevolution alter the ecological dynamics, e.g. changes in population sizes (Ashby et al. 2019; Gutiérrez Al-Khudhairy & Rossberg, 2022), this will impact the supply of *de novo* mutations. The presence of new mutations arising stochastically in the individual reproduction may, in turn, alter the species interactions by regulating the probabilities of host and parasite, or predator and prey, encounters (Frickel et al., 2017). Thus, individual-based models can naturally capture the genetic makeup of populations through mutations (e.g. Huang et al., 2012) and individuals’ responses to ecological and evolutionary mechanisms (Romero-Mujalli et al., 2019). Here, we explored the eco-evolutionary dynamics of a coevolving predator-prey-parasite community using an individual-based model where the parasite is transmitted trophically (Hijar Islas et al., 2024; Figure 1). While demographic fluctuations exist due to species interactions, extinction of species naturally happens depending on the variation in resistance and infectivity shaping host-parasite interactions. We investigated the conditions that promote species coexistence and examined the effects of parasite-mediated reproductive costs on predators and prey on species’ population sizes (i.e., ecological process) and intraspecific diversity (i.e., evolutionary process). Finally, we analysed when super-resistance and super-infectivity can arise in coevolution, and whether the selection of these traits is more determined by population size or by the strength of host-parasite interactions.

**Figure 1.**
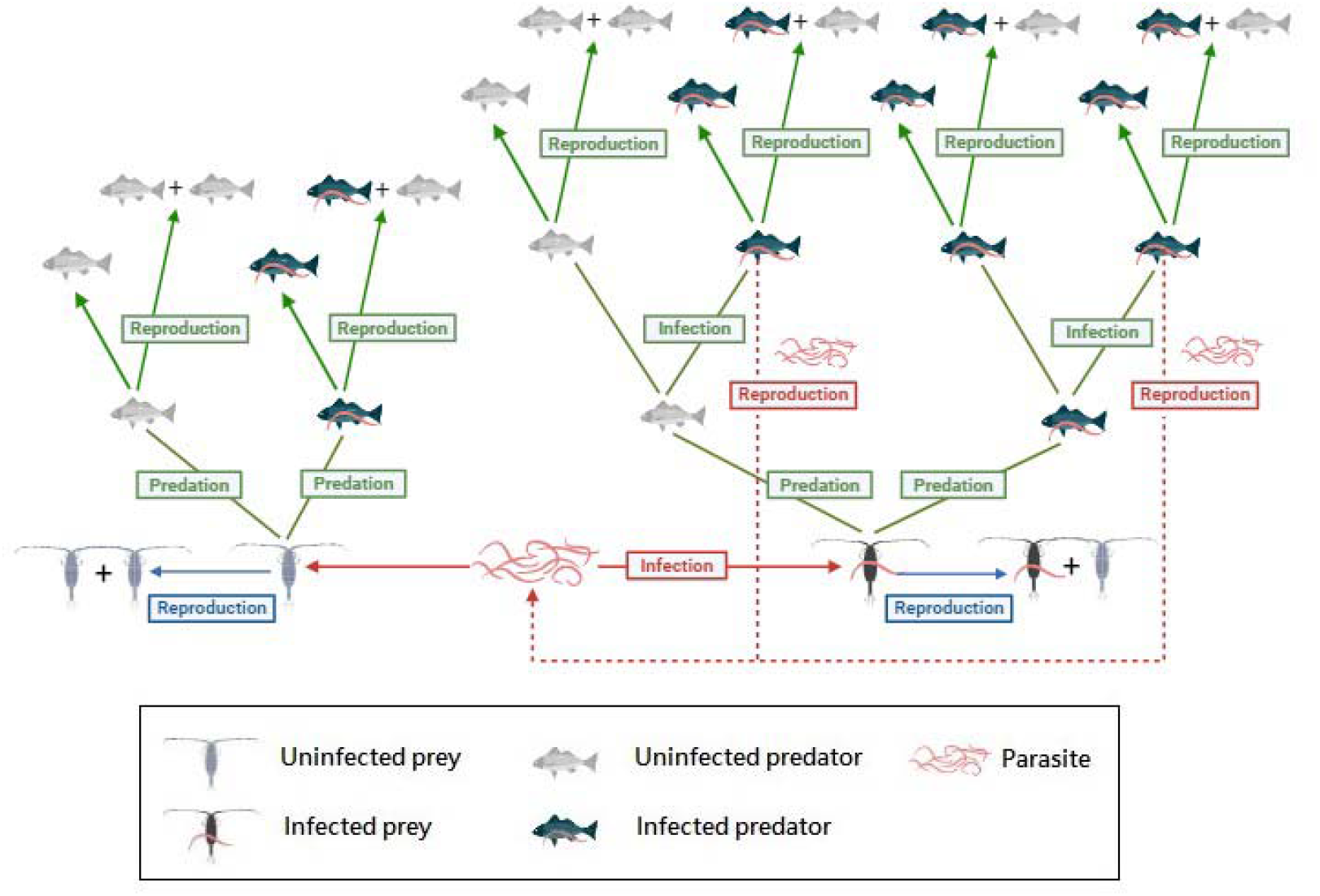
Predator-prey-parasite system. Arrows illustrate all the possible events in the system (red for the parasite, blue for the prey and green for the predator). The parasite can transition from a free-living stage to a state inside the prey (if successfully infecting the prey) until reaching the predator (if successfully infecting the predator), where it reproduces and is released to a free-living stage to begin a new life cycle. The predator is allowed to get infected multiple times.

## Results

We used an individual-based model of a coevolving predator-prey-parasite system in which the parasite is transmitted trophically from the prey to the predator and reproduces when successfully infecting the predator (Hijar Islas et al., 2024). We used a gene-for-gene model for the species interactions (details in Methods), where mutations happen during the reproduction of predator, prey, and parasite species at the rates of *µ*_*y*_, *µ*_*x*_, *µ*_*z*_, respectively. Here, mutation rates are defined per loci per reproduction event. To ensure the three species evolved on similar time scales, we adjusted the mutation rates according to their respective population sizes (Supplementary Note 1). To capture parasite-mediated selection on the prey and the predator, we introduced two parameters, *r*_*x*_ and *r*_*p*_, representing a cost of reproduction inflicted by the parasite on the prey and predator, respectively. The values of *r*_*x*_ and *r*_*p*_ range between 0 and 1, where 0 indicates the parasite completely suppresses the host’s ability to reproduce, and 1 means no impact on the host’s reproduction. We ran multiple independent simulations and investigated how parasite-mediated selection impacts our coevolving system.

### Species coexist more often under weak parasite-mediated selection

Consistent with previous findings on a non-evolving predator-prey-predator system (Hijar Islas et al., 2024), we found that in a coevolving system, the coexistence of all species happens under low parasite-mediated reproductive costs on both prey and predator species (red bars in Figure 2). In the absence of reproductive costs on infected predators (*r*_*p*_ = 1), there is a high likelihood of all species coexistence (97.75% +/-14.83% of the total realisations) in a wide range of reproductive costs on infected prey (from *r*_*x*_ = 0.3 to *r*_*x*_ =1). When the reproductive cost of infected predators is high (*r*_*p*_ = 0.1), we see a significant drop in all species’ coexistence with increasing reproductive costs on infected prey (3.72% +/-18.95% of the total realisations). Due to its complex life cycle, the parasite needs to be transmitted from prey to predators and reproduce in the final host. As, expected, with a complete reproductive suppression of infected predators (*r*_*p*_ = 0), the three species do not coexist.

**Figure 2.**
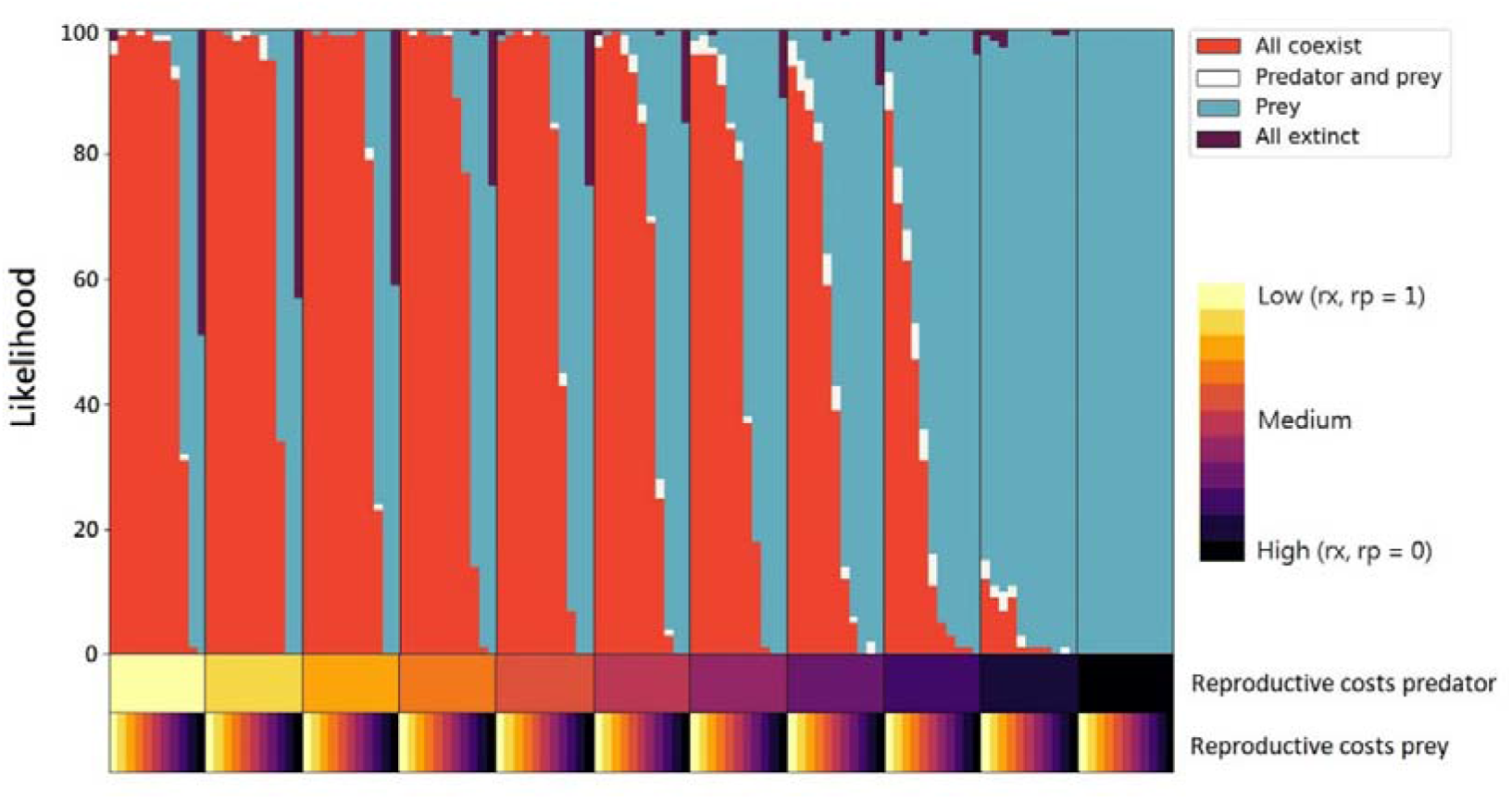
Patterns of coexistence of parasite, prey, and predator. Proportions are taken as a function of parasite-mediated selection on the predator and the prey, from 100 independent realisations. The y-axis shows the number of independent realisations. The x-axis shows the different combinations of parameters ***r***_***x***_ (reproductive costs prey) and ***r***_***p***_ (reproductive costs predator). Note values of ***r***_***x***_ and ***r***_***p***_ are the inverse of reproductive costs: high reproductive cost (marked as darker colours in the x-axis) corresponds to a small value of ***r***_***x***_ and ***r***_***p***_. The values of ***r***_***x***_ and ***r***_***p***_ decrease from left to right in the x-axis from 1 to 0, with an interval of 0.1.

We observe the extinction of the parasite with the coexistence of the predator and prey to be more common with intermediate to low parasite-mediated selection on the predator (white bars in Figure 2). This suggests the emergence of density-dependent transmission of the parasites, which becomes less likely as the predator population size decreases in response to the infection cost. Once the parasite goes extinct, selection for resistance is removed, and the predator population grows again and fluctuates with that of the prey.

With high reproductive costs on infected predators (*r*_*p*_ = 0.1), we show that for a wide range of reproductive costs on infected prey (from *r*_*x*_ =0 to *r*_*x*_ = 1), the predator is likely to go extinct first, followed by the parasite, and the prey population reaches the environment’s carrying capacity (blue bars in Figure 2). On the contrary, with no reproductive costs on infected predators (*r*_*p*_ = 1), we only observe the extinction of both predator and parasite with the prey remaining in a very narrow range, i.e., when reproductive costs on infected prey are relatively high (*r*_*x*_ =0 to *r*_*x*_ = 0.5). This seems counter-intuitive at first sight. Why should the prey remain and reach a high density under a high prey reproduction cost? However, if we combine the observation of all species extinction, high reproductive costs on infected prey also drive all species to extinction (purple bars in Figure 2). Thus, two scenarios are happening when reproductive costs on infected predators are low but high on infected prey. Either the low prey population size leads to its extinction and then the extinction of all species, or it leads to the extinction of the predator first which is followed by that of the parasite, and ultimately a recovery of the prey population.

In addition, the likelihood of all species going extinct increases when the reproductive costs on infected prey remain high but that of infected predators decrease (purple bars in Figure 2). Note, all-species extinction always happens when the prey goes extinct first. These two observations show that prey extinction is a combined result of high parasite-mediated selection as well as predation pressure, especially when parasite-mediated selection is weaker on the predator.

To further understand why and which of the predator, prey and parasite can coexist, we recorded the species’ population sizes and genotypic diversity over time (Figure 3). We examined the relationship between genetic diversity and population size. A lack of correlation could suggest the dominance of a supertype in the system, monopolizing the ecological niche and preventing the emergence and spread of other genotypes (Retel et al., 2019). While most new mutations will fail to invade the coevolutionary system and get lost due to drift, we focused on the effective genotypes making up at least 2% of the population (Figure 3), to avoid the noise due to continuously arising mutations with low frequencies.

**Figure 3.**
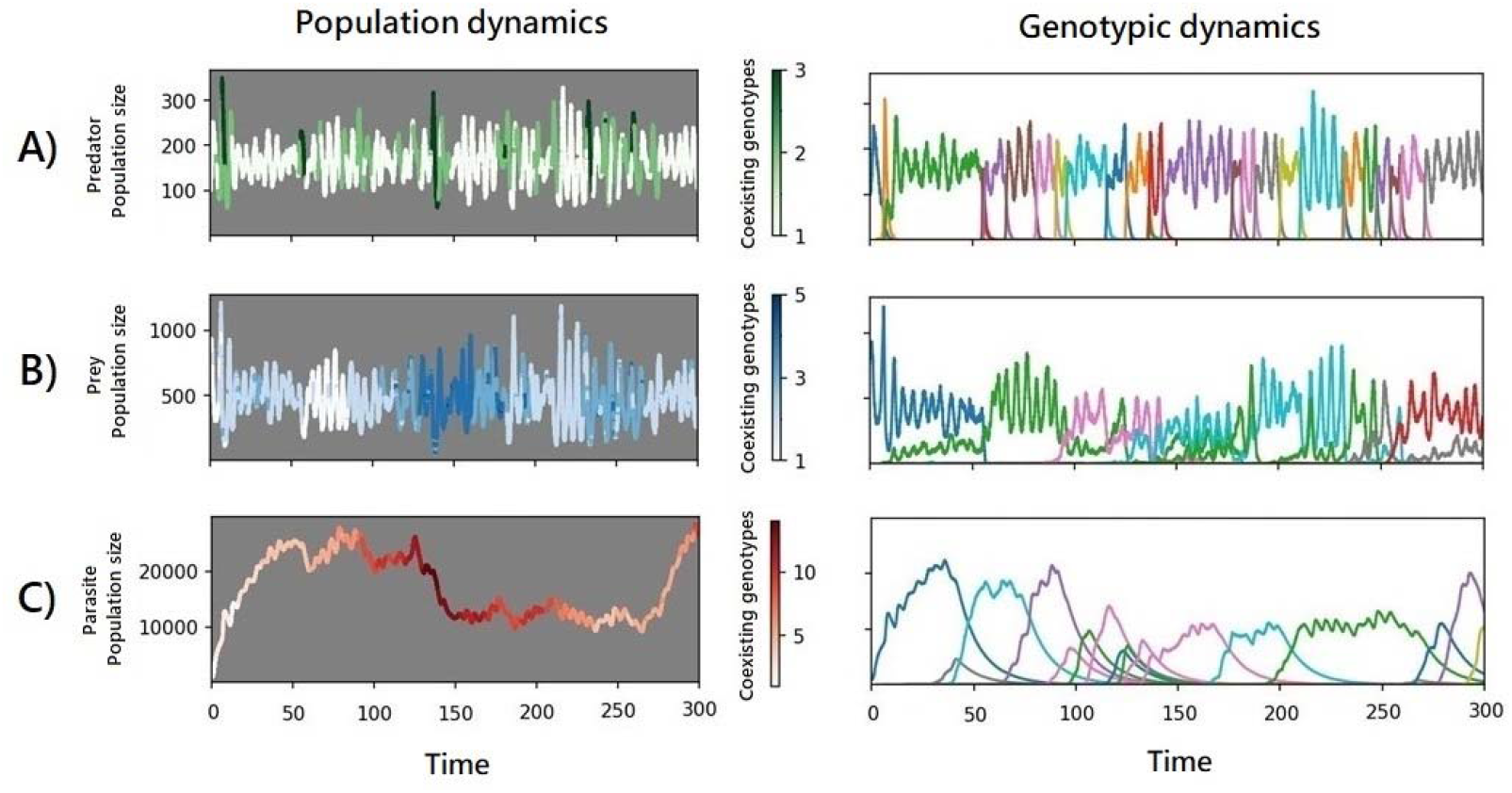
Population and genotypic dynamics from a single realisation. The left panels show the total population dynamics of the species, where darker shading indicates a higher number of genotypes coexisting (see the legend bars on the right side of those panels). The right panels show the abundance of genotypes emerging in each species, where different colours represent different genotypes. A) population and genotypic dynamics of the predator, B) population and genotypic dynamics of the prey, and C) population and genotypic dynamics of the parasite. The parameter set is no parasite-mediated selection on the prey and the predator (i.e., ***r***_***x***_ = 1; ***r***_***p***_ = 1).

### Prey, predator and parasite abundances are maximised in different selection regime

We found the correlations between species’ population sizes were influenced by the strength of parasite-mediated selection on the hosts’ fitness (Table S2). Specifically, in the same parameter space, the parameter sets leading to the highest population size differ for each species (Figure 4A-C). Predators’ population size is highest in the absence of parasite-mediated selection on both the predator and the prey, which maximises reproduction and high prey availability (Figure 4A, upper left corner).

**Figure 4.**
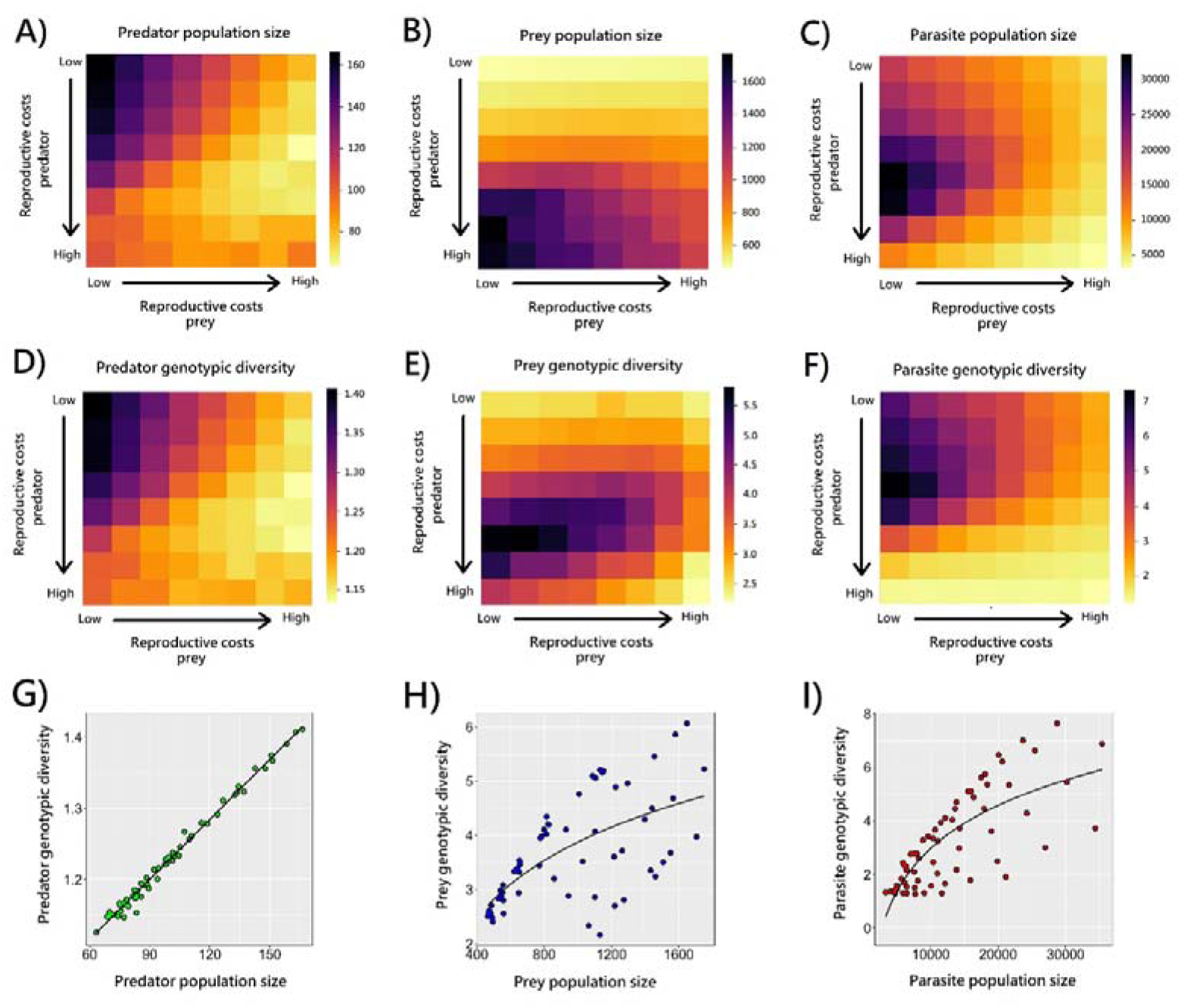
Mean population size and mean genotypic diversity. Genotypic diversity refers to the number of existing genotypes with size >2%. Values are taken as a function of ***r***_***x***_ (reproductive costs prey) and ***r***_***p***_ (reproductive costs predator) when the three species coexist. Note values of ***r***_***x***_ and ***r***_***p***_ are the inverse of reproductive costs: high reproductive cost corresponds to a small value of ***r***_***x***_ and ***r***_***p***_. The mean population size of A) predators; B) prey; and C) parasites; The mean genotypic diversity of D) predators; E) prey; and F) parasites. Each panel displays the average value from 100 independent realisations over a time period of 200-1000 time steps for each realisation. The correlation between mean population size and mean genotypic diversity of G) predators; H) prey; and I) parasites. Each dot displays the average value from 100 independent realisations over a time period of 200-1000 time steps for each realisation.

Prey population size is maximised when parasite-mediated selection on the predator is strong, hence suppressing predator-mediated selection, and when there are no reproductive costs on infected prey (Figure 4B, lower left corner). The magnitude of change in the prey population that results from different strengths of parasite-mediated selection on the prey is small in a parameter space where predators are abundant (Figure S6). This is due to the net effect of both consumers on the prey population: when parasite-mediated selection is absent on the prey, there is more prey available for the predator to consume, leading to a smaller prey population size. Conversely, with increasing parasite-mediated selection on the prey, fewer prey individuals are available for the predator to consume, resulting in weaker predation pressures in a bottom-up control. Interestingly, the mean prey population size reaches its lowest level under no parasite-mediated selection on both hosts (Figure 4B, upper left corner), suggesting the strength of parasite-mediated selection controls the strength of predation pressure when both exist.

Lastly, we found that the parasite population size is maximised with intermediate strength of parasite-mediated selection on the predator combined with no reproductive costs on infected prey (Figure 4C middle left corner). The reduced parasite-mediated selection on the prey maximises uptake by the intermediate host, while the intermediate cost on the final host maximises predator density.

Overall, different conditions favour the maximum population size of each interacting species. Reduced parasite-mediated selection on the prey combined with intermediate levels of selection on the predator, however, not only maintains species coexistence but also maximises population size for all species.

### Drivers of genetic diversity

To understand the link between ecology and evolution, we investigated the correlations between the different population sizes and levels of genetic diversity across replicated realisations. We confirmed that genetic diversity is not merely driven by population sizes, where the highest size always leads to the highest diversity, especially for the prey and parasites (Figures 4B vs 4E, Figures 4C vs 4F).

We observed that the genetic diversity of predators increases with population size (linear model: F_1, 62_ = 6,469; p < 0.0001; Figure 4G). Analysis of predator genotypic dynamics revealed recurrent selective sweeps, consistent with the prediction of an arms race, in which genotypes are unlikely to coexist at relatively high frequencies (Figure 3). With no parasite-mediated selection, the frequency of those selective sweeps is the sole consequence of the mutation rate in large population sizes. When the presence of a parasite imposes strong reproductive costs on the prey population (Figure S7) or the predator population (Figure S8), the size of the predator population decreases, leading to longer and less frequent selective sweeps. In turn, this reduction in predator population size leads to a decrease in the population size of the parasite, resulting in lower infectivity due to a decrease in the mutation supply.

We found that genetic diversity increases with population size in a non-linear manner in both the prey (log-fit model; F_1, 62_ = 44.17; p < 0.0001; Figure 4H) and the parasite (log-fit model; F_1, 62_ = 86.62; p < 0.0001; Figure 4I). Surprisingly, we observed that prey genetic diversity is highest under conditions of intermediate parasite-mediated selection on predators and in the absence of parasite-mediated selection on the prey (Figure 4E). These conditions maximise parasite population size (Figure 4C), rather than prey population size (Figure 4B), suggesting that prey diversity is not a result of the prey population size, but instead the result of prey-parasite coevolution. Also, in this parameter space, the parasite controls the predator’s reproduction rate, allowing the prey to reproduce more rapidly and potentially leading to higher prey genotypic diversity. The absence of strong selection pressures on the prey allows different prey genotypes to coexist, as resistance does not confer any infection-related advantage among genotypes.

We found that in the predator population, the genotypic lifespan is negatively correlated with the population size (log-fit model; F_1, 62_ = 887.1; p < 0.0001; Figure 5A) and with the genotypic diversity (log-fit model; F_1, 62_ = 628; p < 0.0001; Figure 5D). Similarly, in the parasite population, we found that the genotypic lifespan is negatively correlated with the population size (log-fit model; F_1, 62_ = 38.64; p < 0.0001; Figure 5C) and with the genotypic diversity (log-fit model; F_1, 62_ = 261.7; p < 0.0001; Figure 5F). Contrarily, prey genotypic lifespans are longer with medium population size (quadratic regression; F_2, 61_ = 3.931; p = 0.0248; Figure 5B) and with medium genotypic diversity (quadratic regression; F_2, 61_ = 3.049; p = 0.0547; Figure 5E). These trends suggest that strong reciprocal selection pressures between the predator and the parasite lead to the non-coexistence of different genotypes, where advantageous genotypes monopolize the population.

**Figure 5.**
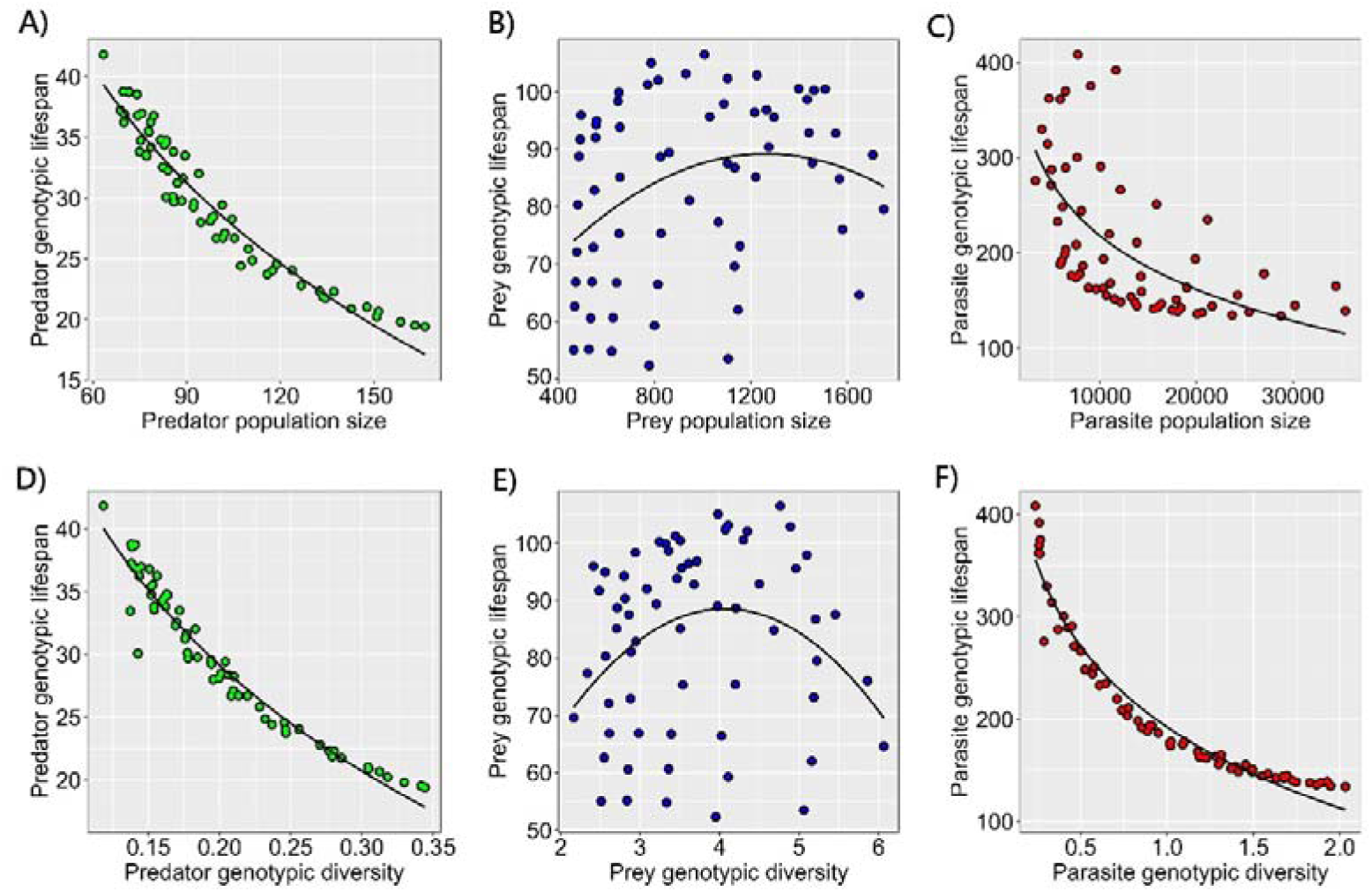
Species’ genotypic lifespan according to the population size and genotypic diversity. Genotypic lifespan refers to the number of time steps from the moment a genotype emerged until it went extinct. Genotypic diversity refers to the number of existing genotypes with size >2%. Values are taken as a function of the strength of parasite-mediated selection on the predator and the prey when the three species coexist. Correlations between mean population size and mean genotypic lifespan of the A) predator, B) prey, and C) parasite. Correlations between mean genotypic diversity and mean genotypic lifespan of the D) predator, E) prey, and F) parasite. Each dot displays the average value from 100 independent realisations over a time period of 200-1000 time steps for each realisation.

While genotypic dynamics are also shaped by selective sweeps, the lifespans of the parasite genotypes are longer than those of the prey and predators, particularly under high reproductive costs on the predator that result in low parasite population size and low genotypic diversity (Figure S8). As the parasite needs to reach the predator to reproduce, when it imposes high reproductive costs on infected predators, the parasite population size decreases, resulting in fewer mutations. If the predator evolves resistance, however, the parasite population persists with low infectivity and coexists with the predator (Figure S8).

### Evolution of super-resistant and super-infective genotypes

Our gene-for-gene coevolving model enables us to detect the emergence of super-resistant prey and/or predator genotypes, as well as super-infective parasite genotypes. A super-resistant host is a genotype that has resistant alleles in all modelled loci, while a super-infective parasite is a genotype that has infective alleles in all modelled loci. Our results suggest that the evolution of these supertypes in a focal species is mediated by the population size of interacting species, rather than by its own population size (Figure 6). For instance, when all species coexist, predators are more likely to evolve super-resistant genotypes under conditions of intermediate parasite-mediated selection on the predator combined with no reproductive costs on the prey (11% of the realisations, Figure 6A), which is the parameter space with highest parasite population size (Figure 4C) and strongest selection pressures on the predator. The evolution of super-resistant prey genotypes is mostly driven by predator population size rather than parasite-mediated selection on the prey. This suggests that parasite pressure drives the evolution of prey resistance through the predator, leaving a signature of indirect parasite-mediated selection. Prey are more likely to evolve super-resistant genotypes with intermediate reproductive costs on the predator and the prey due to lower predation pressure (22% of the realisations; Figure 6B). Parasites on the other hand are more likely to evolve super-infectivity under no parasite-mediated selection on the prey, particularly combined with intermediate reproductive costs on the predator (30% of the realisations; Figure 6C). This suggests that the probability of the evolution of super-infectivity scales with parasite population size, and super-infective parasite genotypes are selected rapidly when the reproductive costs on both hosts are not high.

**Figure 6.**
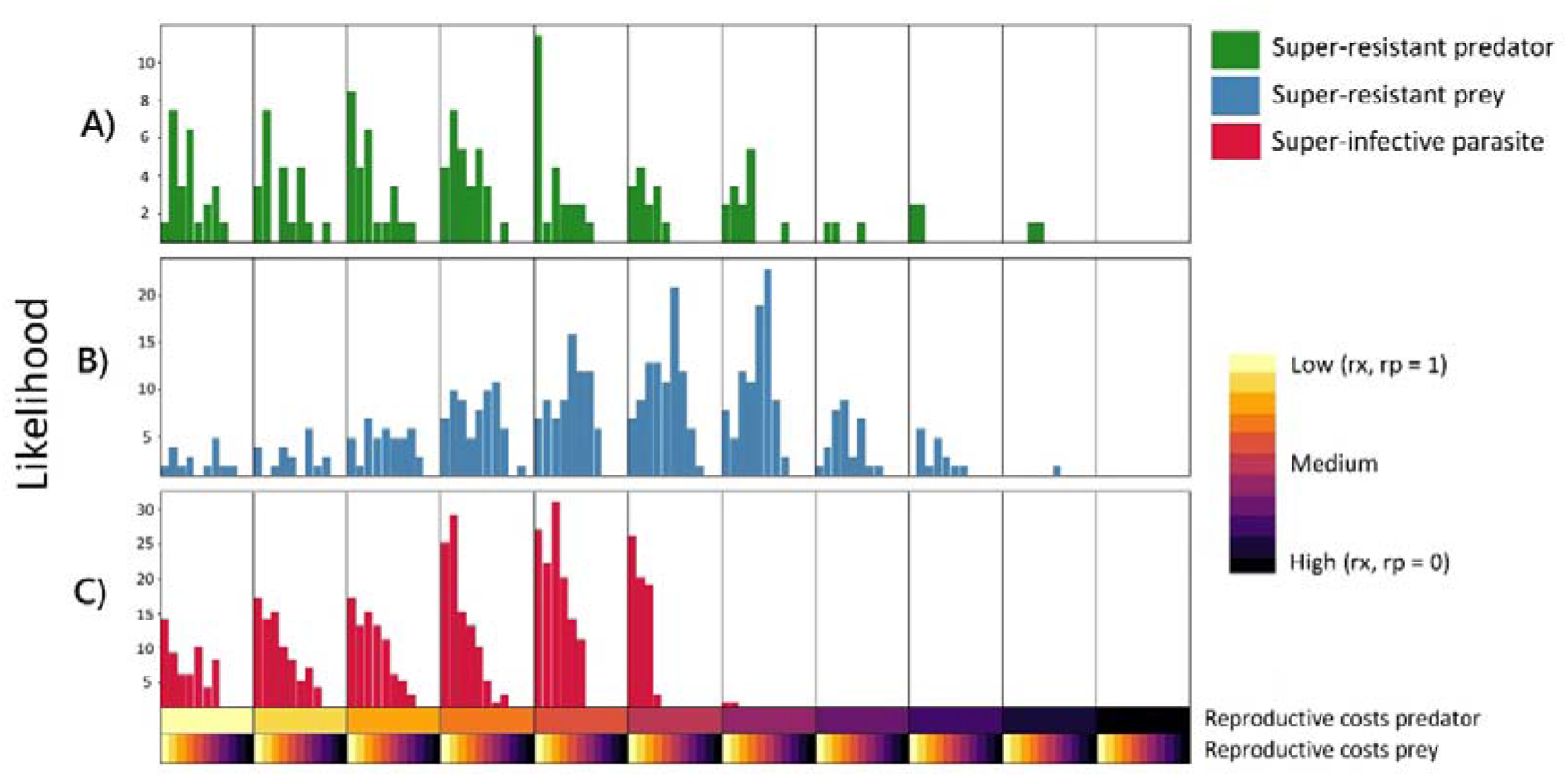
The emergence of supertypes when the three species coexist. A) Evolution of a super-resistant predator genotype; B) Evolution of a super-resistant prey genotype, and C) Evolution of a super-infective parasite genotype. Proportions are taken as a function of the strength of parasite-mediated selection on the predator and the prey, from 100 independent realisations (shown on the y-axis). The y-axis shows the number of independent realisations. The x-axis shows the different combinations of parameters ***r***_***x***_ (reproductive costs prey) and ***r***_***p***_ (reproductive costs predator). Note values of ***r***_***x***_ and ***r***_***p***_ are the inverse of reproductive costs: high reproductive cost (marked as darker colours in the x-axis) corresponds to a small value of ***r***_***x***_ and ***r***_***p***_. The values of ***r***_***x***_ and ***r***_***p***_ decrease from left to right in the x-axis from 1 to 0, with an interval of 0.1.

## Discussion

While it is well known that the dynamic interplay of ecological and evolutionary processes shapes species interactions, the role of these eco-evolutionary dynamics in complex systems such as food webs remains to be generalised. By integrating host-parasite and predator-prey interactions in a coevolving system where mutations can emerge, we demonstrate that species coexistence is more likely under low parasite-mediated selection on both host species. Population sizes and genotypic diversity increase with intermediate parasite-mediated selection (i.e., reproductive costs) on the predator combined with no parasite-mediated selection on the prey. Our results show that consumer-resource correlations are determined by the strength of parasite-mediated selection and the hosts’ fitness. We also show that multiple interspecific interactions drive the emergence of supertypes, in which host and parasite fitness is maximised. Overall, our findings suggest that species coexistence and community diversity are mediated by host-parasite coevolution, which is shaped by multispecies ecological interactions.

We show that parasite-mediated selection determines species’ coexistence (Figure 2). In our coevolving predator-prey-parasite system, the prey is a shared resource for parasites and predators. Due to this niche overlap, the prey goes extinct under strong parasite and predator-mediated selections, ultimately leading to the extinction of all three species. The predator is a consumer of the prey and a resource of the parasite, so a large predator population size is beneficial for the parasite but detrimental to the prey. With few prey individuals, the predator population collapses, followed by the parasite population. The parasite remains if there is enough prey for the predator to consume and enough predators to act as final hosts. Under strong parasite-mediated selection, the evolution of resistance rescues the system and allows species coexistence (Figure S8). This theoretical prediction matches the experimental demonstration of the rescue mechanism detected in virus-host-predator systems, where host-virus coevolution facilitates the coexistence of competing consumers (Frickel et al., 2017).

In bacterial-phage systems, phages evolve faster than their bacterial host (Campbell, 1961; Heilmann et al., 2010; Rudolf & Antonovics, 2005). Due to their rapid evolution, phages can coexist with their bacterial host through ecological mechanisms such as reduced phage dispersal (Brockhurst et al., 2006), and density-dependent spatial bacteria refuges (Heilmann et al., 2012). In our system, parasite infectivity evolves at the same pace as host resistance because of the parasite’s complex lifecycle. The evolution of resistance in the predator can be advantageous for the parasite under strong parasite-mediated selection on the predator, and the three species can coexist with the predator and parasite remaining at small population sizes (Figure S8).

When integrating parasites in food webs, predator-prey correlations depend on the strength of parasite-mediated selection, which determines predator and prey population sizes as well as genotypic diversity. Parasites alter predator-prey interactions via predator-mediated effects (top-down cascade) on the prey and prey-mediated effects (bottom-up effect) on the predator (Anaya-Rojas et al., 2019). The availability of both hosts is highest under no parasite-mediated selection on the prey and with intermediate reproductive costs on the predator given that the predator is a resource to the parasite and a consumer to the prey (Figures 4C & 4F). This parameter space maximises the population size of the three species. This prediction matches observation where parasites that use multiple hosts to complete their life cycles often impose relatively no harm to the intermediate host and are very harmful to the final host in which they reproduce (De Castro & Bolker, 2005; Scholz et al., 2021). Interestingly, we find that the prey is more abundant under strong parasite-mediated selection on both hosts (Figure 4B, lower right corner) than under no parasite-mediated selection (Figure 4B, upper left corner). This suggests that parasites control prey population size via indirect effects on the predator population and that the maintenance of genetic diversity is driven by multiple selective agents.

In conditions where selection pressures are strong, a genotype may dominate the ecological niche due to evolutionary-mediated priority effects (De Meester et al., 2016). Contrarily, when selection pressures are weak, more genotypes are likely to co-occur and drift neutrally (De Meester et al., 2016). In our system, genotypic diversity increases in the absence of parasite-mediated selection on the hosts because the evolution of resistance alleles confers no fitness advantage. On the other hand, the lifespan of a genotype reflects its dynamics and the sweeps it undergoes when intraspecific diversity is low. The lifespans of the genotypes are determined by the population size because the mutation supply increases with population size and provides more fuel for adaptation (Retel et al., 2019). Even under weak parasite-mediated selection, host-parasite interactions can lead to rapid allele frequency shifts, even those initially rare (Gokhale et al., 2013), as demonstrated experimentally in three-spined stickleback for instance (Eizaguirre et al., 2012).

We show that host and parasite genotypic dynamics are shaped by consecutive selective sweeps, where the lifespans of the parasite genotypes are longer than those of the prey and predators (Figures 3, S7 & S8). These genotypic dynamics have been shown experimentally in a host-virus system (Retel et al., 2019). Interestingly, we see that the predator genotypic dynamics are governed by the succession of many selective sweeps rather than coexisting genotypes, even in the absence of parasite-mediated selection (Figure 3). This suggests that predator-prey interactions alone can facilitate the rapid emergence of resistance genotypes. Note that since we use a haploid genetic system, mutations are directly exposed to selection. The fitness advantage of those mutations depends on the allelic states of co-occurring host-parasite genotypes. Our results suggest that recurrent selective sweeps are common in the predator population and shape long-lasting coevolutionary dynamics with the parasite.

In host-parasite interactions, population and allele dynamics interact to determine coevolution (Retel et al., 2019). Our results show that host-parasite coevolution is influenced by other species interactions. For instance, both predator and parasite-mediated selections drive the evolution of super-resistant prey genotypes, but mostly the predator. Indeed, we show the prey is more likely to evolve super-resistant genotypes under relaxed predation pressure that leads to large population size, hence increasing the mutation supply. Under those conditions, the presence of parasites selects for resistant genotypes in the prey population (Figure 6B). This finding is consistent with previous experimental work (Frickel et al., 2017; Brunner et al., 2017) showing that parasite-mediated predation drives the evolution of the prey. Super-infective parasite genotypes evolve under low parasite-mediated selection on both hosts because high infectivity leads to host extinction when the parasite decreases largely its host fitness (Figure 6C). The parasite is more likely to evolve super-infectivity under a large population size, which is exactly the parameter space where the availability of both hosts increases. As the predator is more likely to evolve super-resistant genotypes when parasites are abundant (Figure 6A), the parasite needs to counter-evolve infectivity to persist and complete its lifecycle.

We know that host-parasite coevolution is determined by the population dynamics and genetic makeup of interacting species (Buckingham & Ashby, 2022). Here, we demonstrate that, in a complex system, host-parasite coevolution is also shaped by interspecific interactions, where prey-parasite interactions are altered via top-down control by the predator, and predator-parasite interactions are mediated by the prey in a bottom-up manner. Host-parasite coevolution, in turn, shapes population dynamics, as the evolution of resistance mitigates the detrimental effects on the hosts. The changes in predator and/or prey population size that result from parasite-mediated selection, hence, are associated with evolution-mediated trophic cascades that impact the whole community’s abundance and diversity. This has been demonstrated experimentally, in a non-microbial system, where the evolution of resistance in predator fish impacts prey community structure (Brunner et al., 2017).

This dynamic interplay of ecology and evolution becomes evident at different strengths of parasite-mediated selection on the hosts. The degree of these effects determines the magnitude of change in population sizes which, in turn, determines the evolution rates for resistance/infectivity shaping host-parasite interactions.

## Methods

### Model of host-parasite coevolution

We constructed the host-parasite coevolution by extending a multi-locus gene-for-gene model (Sasaki, 2000). We consider a prey (i.e., intermediate host) genotype for resistance *j* = *j*^1^, *j*^2^ *… j* ^*n*^, a predator (i.e., final host) genotype for resistance *l* = *l*^*1*^, *l*^*2*^*… l* ^*n*^, and a trophically transmitted parasite genotype for infectivity *k* = *k*^1^,*k*^2^*… k* ^*n*^, where *n* corresponds to the number of loci (i.e., genotype length). For different genotypes, the allelic state of prey resistance, predator resistance, or parasite infectivity, at each locus of the genotypes is a binary number, where ‘0’ corresponds to susceptible/non-infective alleles and ‘1’ to resistant/infective alleles. These allelic states are reversible, where mutations can lead to allelic changes in both directions for all the genotypes (i.e., from ‘0’ to ‘1’, or from ‘1’ to ‘0’). *X* ^*j*^ is the prey of genotype *j, Y*^*l*^ is the predator of genotype, and z ^k^ is the parasite of genotype *k*, where *j, l* and *k* are vectors of 2 ^*n*^ possible combinations.

For simplicity, we focus on haploid and asexual hosts and parasites. The parasite genotypes possess 10 loci for infective/non-infective alleles, and both the prey and predator hosts genotypes possess 10 loci for resistant/susceptible alleles. Random mutations arise in the three species after a reproduction event with rates *µ*_*x*_ for the prey, *µ*_*y*_ for the predator, and *µ*_*z*_ for the parasite. *µ*_*x*_, *µ*_*y*_, *µ*_*z*_, are values between 0 and 1, i.e., the mutation probability per locus per reproduction. To simulate the mutation event of any genotype per reproduction, we draw a random number between 0 and 1 for each locus. If any of the n random numbers are smaller than the mutation probability, a mutation happens, otherwise, the genotype is the same as the parent.

All the hosts (predator and prey) are initially susceptible, and all parasites are initially non-infective except to the initial hosts (i.e., all genotypes are initially simulated as strings of ‘0’). If the genotype of the initial prey *X* ^*j*^ is 00…00, and the genotype of the initial parasite *z* ^*k*^ is 00…00, then the initial effective resistance is *r*(*X* ^*j*^, *Z* ^*k*^) = 0, and the probability of successful infection of the initial parasite to the initial prey is *Q*_*x*_ (*x* ^*j*^, *z* ^*k*^) = 1. Likewise, if the genotype of the initial predator *Y*^*l*^ is 00…00, and the genotype of the initial parasite *Z* ^*k*^ is 00…00, then the initial effective resistance is *r*(*Y*^*l*^, *Z* ^*k*^) = 0 and the probability of successful infection of the initial parasite to the initial predator is *Q*_*y*_ (*Y*^*l*^, *Z* ^*k*^) = 1 (see details of the probability of successful infection in Supplementary Note 2).

We consider well-mixed populations, where the encounter rates between the hosts and parasites are density-dependent. Ultimately, the probability of successful infection between a parasite and a host is a function of the allelic states across all loci of the parasite and the host genotypes. We consider each position in the host genotype to see whether it is resistant to a particular locus of the interacting parasite. The concrete value of host resistance r is the number of resistant alleles in the host genotype (‘1’) matching the same position as non-infective alleles in the respective parasite genotype (‘0’).

### The individual-based model

We used an individual-based model of a predator-prey-parasite system, where the parasite is transmitted trophically from infected prey to predators (Hijar Islas et al., 2024). We allowed a coevolving predator-prey-parasite community, where mutations can happen in the prey, predator, and parasite genotypes in reproduction. The allelic states across all loci of the predator, prey and parasite genotypes ultimately determine the probability of successful infection. Each individual-level event occurs with a type-specific rate indicated by the parameters above the arrows in our reactions. The model parameters are shown in Table S1 and the choice of parameter values is shown in Supplementary Note 1. While reactions and their rates are defined at the individual level, how often e.g., reproduction of prey happens at the population level will also depend on the population size.

### Possible events only in the prey population

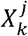 is a prey of genotype *j*. When *k* = 0, the prey does not carry a parasite. Suppose such a prey reproduces an offspring 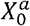. The new offspring has the same genotype as the parent when *a* = *j*, and is a mutant when *a* ≠ *j*. The probability of no mutation happening during a reproduction event is (1 - *µ*_*x*_) ^*n*^, where *µ*_x_ ∈ [0,1] is the probability of a mutation per locus and n is the number of loci in the genotype. We note here that the prey offspring is parasite-free and can be infected by the free living-parasite population in the environment.

In the absence of the other species the prey populations can be described by the following reactions, where *g*_*x*_ ∈ *R* ^+^ is the reproduction rate of the prey, *r*_*x*_ ∈ [0,1] is the reduction of an infected prey reproduction, *K* ∈ *R* ^+^ is the environmental carrying capacity of the prey population, and *d*_*x*_ ∈ *R* ^+^ is the intrinsic death rate.

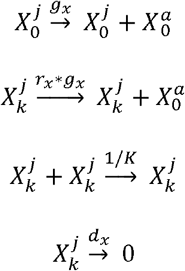

While we already implement a cost of infection in reproduction, we assume that infection does not influence the competition outcome nor intrinsic death for the sake of simplicity. However, this cost of death can be easily implemented if necessary.

### Possible events in the parasite population

*Z^k^* is a free-living parasite of genotype *k*. 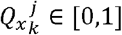 is the probability of successful infection of a parasite with genotype to prey with genotype j, which depends on the allelic states of the interacting prey-parasite genotypes. *S* ∈ *R* ^+^ is the encounter rate between prey and free-living parasites. When a parasite successfully infects the prey, the parasite is removed from the free-living parasite population. Otherwise, it remains in the free-living parasite population. We note here that the parasite does not reproduce in the prey. As it is computationally expensive to simulate single prey individuals carrying multiple parasite genotypes, we assume that the free-living parasites only infect uninfected prey.

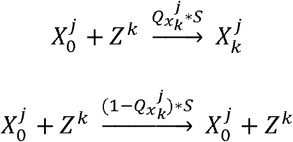

All parasite individuals have an intrinsic death rate *d*_*z*_ ∈ *R* ^+^, leading to the ultimate extinction of the parasite if there are no predator and prey species.

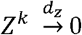

### Possible events in predator population

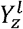 is a predator of genotype *l*. When z = 0, the predator does not carry a parasite. Let us introduce the notation 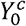, where c = *l* when the new offspring is the same genotype as the parent, and c ≠ *l* when the new offspring is a mutant. The probability of no mutation happening during reproduction is (1 − *µ*_*y*_) ^*n*^, where *µ*_*y*_ ∈ [0,1] is the probability of a mutation per locus and n is the number of loci in the genotype. The predation rate per individual encounter between a prey and a predator is *f*_*y*_. The probability of a reproduction event occurring given a predation event is *k*_*y*_ ∈ [0,1]. We note here that the predator offspring is parasite-free, 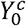. When an uninfected prey meets an uninfected predator, we have the following reactions.

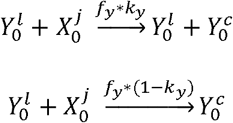

When a prey of genotype *j*, infected with a parasite of genotype meets an uninfected predator of genotype *l*, the predator can be infected with a probability 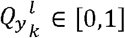. If the predator is successfully infected by the parasite through the consumption of infected prey, the infected predator’s ability to reproduce is affected by the presence of the parasite and hence is reduced according to the parameter *r*_*p*_ ∈ [0,1]. Meanwhile, the parasite transmitted from the primary host, the prey, will reproduce within the definitive host, the predator, and have *nZ* ^*b*^ ∈ z ^+^ offspring per successful infection. The original parasite stays in the predator and later dies together with the predator stochastically. The parasite offspring, *nZ* ^*b*^, are released to begin a new life cycle, where *b* =*k* when the new offspring is the same genotype as the parent, and *b ≠ k* when the new offspring is a mutant. The probability of no mutation happening during reproduction is (1-*µ_z_*)^*n*^, where *µ_z_* ∈ [0,1] is the probability of a mutation per locus and n is the number of loci in the genotype.

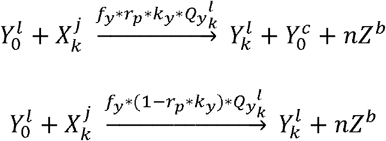

If the parasite is not successful in infecting the predator through trophic transmission, the prey and parasite, 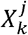, die. In this case, there is still a reduction in predator reproduction due to exposure *r_e_* ∈ [0,1] because it triggers an immune response, which is energetically costly, even in the absence of infection (Eizaguirre et al., 2012). We assume that the reproductive cost due to infection is larger than the one due to exposure. Hence, *r_p_ < r_e_* ≤ 1.

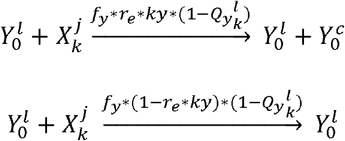

When an infected predator 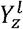 (where *z* refers to any parasite genotype) consumes a parasite-free prey of genotype *j*, the infected predator can reproduce at the cost *r*_*p*_. Note parasite reproduction already happened when this predator was first time infected, thus is not redundantly included here.

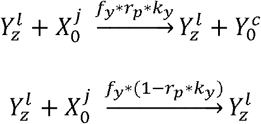

Finally, we consider the remaining possible reactions where an infected prey meets an infected predator. When an infected predator 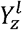 consumes a prey of genotype *j* infected with a parasite of genotype *k*, the predator can produce offspring without the parasite 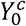. If the infection fails, the prey and parasite, 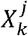, die. We apply the reproductive cost of the predator as before by parameter *r*_*e*_.

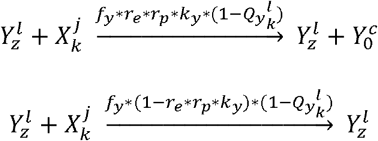

If the infected predator is successfully infected again by the newly encountered infected prey, its ability to reproduce is further reduced by the same parameter *r_p_*. The parasite transmitted by this newly encountered prey will reproduce in its final host because this is the first time this predator is infected by this individual parasite. As it is computationally expensive to simulate multiple genotypes in single predator individuals, and we do not need to keep track of those genotypes as they are no longer transferred to other hosts, we simply use the notation 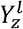 to refer to the infected predator after a new infection, where *z* refers to any parasite genotype.

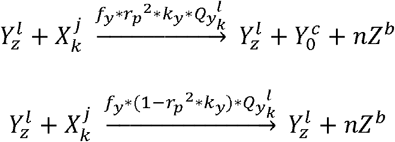

All predator individuals have an intrinsic death rate *d*_*y*_ ∈ *R* ^+^, leading to the ultimate extinction of the predator if there are no prey species. While we already implement a cost of infection in reproduction, we assume that infection does not influence intrinsic death for the sake of simplicity. However, this cost of death can be easily implemented if necessary.

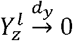

### Model Simulations

We ran multiple independent realisations using the Gillespie algorithm (Gillespie, 1977; Pineda-Krch, 2008) to investigate how parasite-mediated selection impacts our coevolving system. We used different combinations of the parameters *r_x_* and *r_p_*, to capture the parasite-mediated selection on the prey and the predator, respectively. Note that the parameters *r_x_* and *r_p_* range between 0 and 1, with an increment of 0.1, i.e., *r*_*x*_ = 0, 0.1, 0.2, …, 1, and *r*_*p*_ = 0, 0.1, 0.2, …, 1, where 0 indicates the parasite completely suppresses the host’s ability to reproduce, and a value of 1 that there is no parasite impact on the host reproduction (see details in the individual-based model). We ran 100 independent realisations for each combination of *r*_*x*_ and *r*_*p*_. In every realisation, we used the same initial population sizes, i.e., 800 prey individuals, 1,000 parasite individuals, and 100 predator individuals (Hijar Islas et al., 2024). See the choice of the number of realisations in Supplementary Note 3.

## Data Analyses

To obtain the likelihood of species coexistence, we performed 100 independent realisations for each parameter combination *r*_*x*_ and *r*_*p*_. We recorded the number of realisations in which the predator, prey, or parasite went extinct.

The population size and genotypic diversity of each species were recorded in single time points when the three species coexisted, for each parameter combination of *r*_*x*_ and *r*_*p*_. To account for the population size and genotypic diversity in the prey and predator, we considered infected and non-infected individuals. To account for the population size and genotypic diversity in the parasite, we considered the free-living individuals. To avoid the over-representation of initial states where the species have not co-evolved away from the initial conditions, we used the average of 100 realisations from time point 200-1000 to represent the population size and genotypic diversity in each parameter combination of *r*_*x*_ and *r*_*p*_. Supplementary Note 3 explains the choice of the time points for averages of population sizes and genotypic diversity.

We performed a linear model to explore the correlation between the population size and the genotypic diversity of the predator. We conducted fit logarithmic models to explore the correlations between the population size and the genotypic diversity of the prey and the parasite. We additionally conducted fit logarithmic models to explore the correlations between the population size and genotypic lifespan, as well as between the genotypic diversity and genotypic lifespan of the predator and the parasite. Finally, we performed a quadratic regression to explore the correlations between the population size and genotypic lifespan, as well as between the genotypic diversity and genotypic lifespan of the prey. We refer to genotypic lifespan as the number of time steps from the moment a genotype emerged until it went extinct. To avoid the noise due to continuously arising mutations with low frequencies, we focused on the effective genotypes making up at least 2% of the population.

Finally, for the likelihood of the emergence of supertypes, we recorded the number of realisations in which a supertype emerged when the three species coexisted.

