## Supplementary Information. for "Complex life cycles and parasite-mediated trophic cascades drive species coexistence and the maintenance of genetic diversity"

**Contents**

**Supplementary note 1.** Choice of parameter values

**Supplementary note 2.** Probability of successful infection

**Supplementary note 3.** Choice of time points for calculating the mean population size and genotypic diversity of each entity

**Supplementary Figures**

**Supplementary Tables**

**List of Figures**

**Figure S1.** Parameterization of the model for species coexistence at equilibrium.

**Figure S2.** Population and genotypic dynamics of predator, prey, and parasite, according to different mutation rates of the parasite.

**Figure S3.** Probabilities of successful infection according to genotypic changes with *de novo* mutations.

**Figure S4.** Exponential distribution of probabilities of successful infection according to increasing values of σ.

**Figure S5.** Mean population size and genotypic diversity from the average of 100 independent realisations.

**Figure S6.** Equilibrium points of prey population dynamics under different selection pressures on the prey and predator.

**Figure S7.** Population and genotypic dynamics under high reproductive costs on the prey and low reproductive costs on the predator, from a single realisation.

**Figure S8.** Population and genotypic dynamics under low reproductive costs on the prey and high reproductive costs on the predator, from a single realisation.

**List of Tables**

**Table S1.** Model parameters.

**Table S2.** Correlations between mean population size of predators, prey, and parasites as a function of parasite-mediated selection on the prey and the predator.

**Supplementary Note 1: Choice of parameter values**

While we are interested in a coevolving system, we parameterized the model where species can stably coexist at equilibrium of our corresponding deterministic dynamics (Table S1; Híjar-Islas et al., 2023), in the absence of fitness costs on the prey and predator ($r_{x}=1$ and $r_{p}=1$, respectively) and with fixed probabilities of successful infection on the predator and the prey ($Q_{y}=1$ and $Q_{x}=1$, respectively).

For all simulations, we used the same initial population sizes, i.e., 800 prey individuals, 1,000 parasite individuals, and 100 predator individuals. Note that changes in the initial population sizes do not change the population sizes at equilibrium but the amplitude of the oscillations (Figure S1). The amplitudes of the underlying dynamics are a natural ecological factor in stochastic dynamics, where species can go extinct if the population size cycles into a small population size. On the contrary in a deterministic system with only equations, species will always recover to their equilibrium. Thus, besides using deterministic systems to give insight into the average behaviour, it is important to implement the stochastic simulations of an evolving system in species interactions.

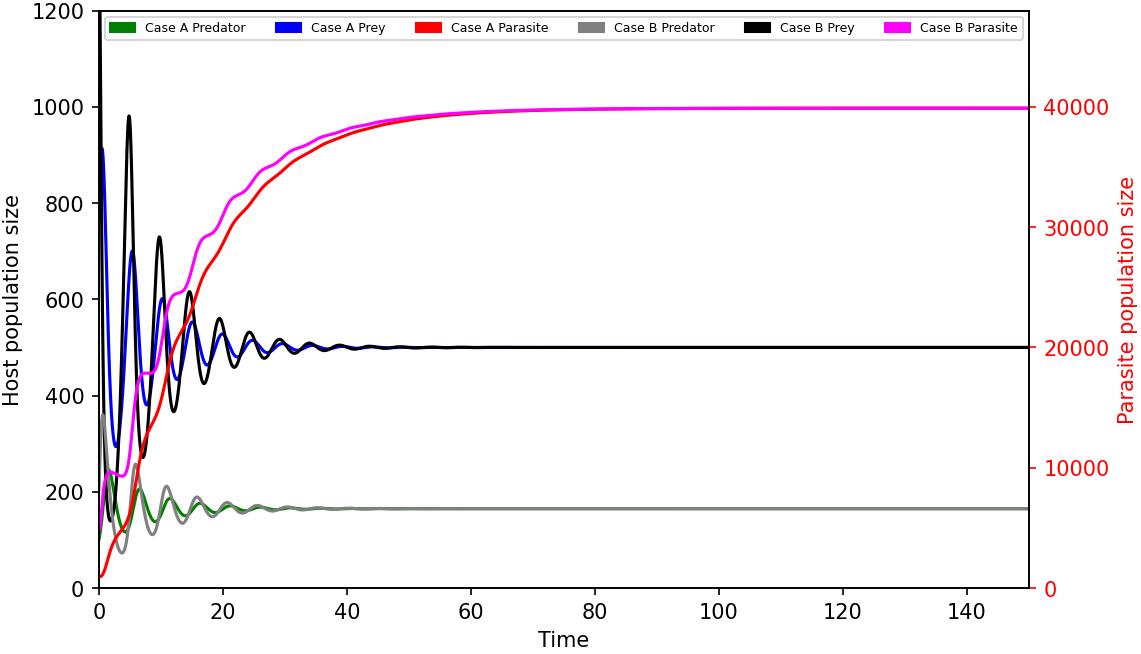

**Figure S1. Parameterization of the model for species coexistence at equilibrium.** Parameter values were explored in the absence of fitness costs on the prey and predator ($r_{x}=1$ and $r_{p}=1$, respectively) and with fixed probabilities of successful infection of the predator and the prey ($Q_{y}=1$**,** $Q_{x}=1$**,** respectively). Case A (800 prey individuals, 1,000 parasite individuals, and 100 predator individuals) and Case B (1500 prey individuals, 5000 parasite individuals, and 200 predator individuals) indicate two different initial population sizes and converge to the same equilibrium because all parameters are the same. Parameter values were chosen following Híjar-Islas et al., 2023.

**Mutation rates**

In real biological systems, mutation rates vary in different species, depending on multiple factors such as genome size, presence, or absence of mutators, or environments. We chose mutation rates of predator, prey, and parasite here, such as the three species can evolve on a similar time scale and allow the emergence of multiple genotypes for each entity under a reasonable computational power. The effective mutation rate is described as $N\times{n\times\mu}_{u}$ where $N$ is the population size, $n$ is the number of loci, and $\mu_{u}$is the mutation rate per locus per reproduction ($\mu_{x}$ for the prey, $\mu_{y}$ for the predator, and $\mu_{z}$ for the parasite). A reasonable value for the effective mutation rate, $N\times{n\times\mu}_{u}$, is 0.1. From the populations’ equilibrium points shown in Figure S1, the mutation rate per locus per reproduction of the prey and predator ($\mu_{x}$ and $\mu_{y}$, respectively) are as follows:

$\mu_{x}$ = $\frac{effective mutation rate}{N\times n}$ = $\frac{0.1}{500\times10}$ = 2$\times$10^-5^

$\mu_{y}$ = $\frac{effective mutation rate}{N\times n}= \frac{0.1}{200\times10}$ = 5$\times$10^-5^

Due to the parasite’s complex life cycle, the parasite needs to infect both hosts to reproduce. This initial delay in the evolution of infectivity leads to a decrease in the parasite population size, and, consequently, the effective mutation rate (Figure S2A). We tuned the value of $\mu_{z}$, to ensure that the evolution of parasite infectivity happened on a similar time scale as the predator and prey, in initial conditions (Figure S2B).

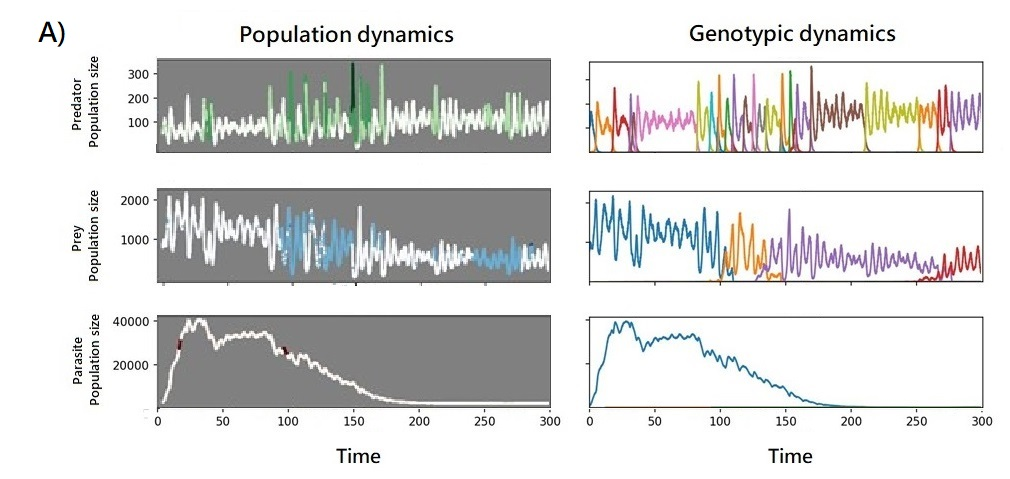

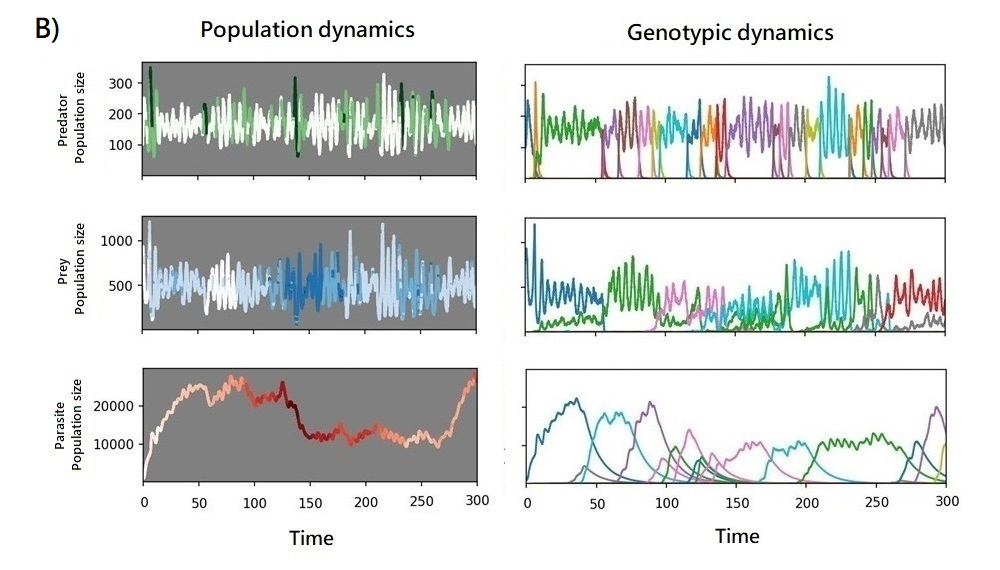

**Figure S2. Population and genotypic dynamics of predator, prey, and parasite, according to different mutation rates of the parasite.** The Figure shows the dynamics from a single realisation, in the absence of reproductive costs on the prey and the predator ($r_{x}=1$ and $r_{p}=1$, respectively). A) $\mu_{z}$ = 2.5x10^-7^. B) $\mu_{z}$ = 6x10^-6^.

**Table S1. Model parameters.** Parameter values were chosen following Híjar-Islas et al., 2023.

| **Parameter** | **Definition** | **Value** |
| --- | --- | --- |
| $g_{x}$ | Reproduction rate of the prey | 2 |
| $r_{x}$ | Reproductive cost on the prey due to parasite infection | [0-1] |
| $d_{x}$ | Intrinsic death rate of the prey | 0.1 |
| $K$ | Environmental carrying capacity of the prey | 2000 |
| $\mu_{x}$ | Mutation rate of the prey (per locus per reproduction) | 2x10^-5^ |
| $S$ | Scaling factor for prey-parasite interactions | 5x10^-4^ |
| $nZ$ | Offspring per reproduction of the parasite | 6 |
| $d_{z}$ | Intrinsic death rate of the parasite | 0.09 |
| $\mu_{z}$ | Mutation rate of the parasite (per locus per reproduction) | 6x10^-6^ |
| $f_{y}$ | Predation rate | 0.01 |
| $k_{y}$ | Reproduction rate of the predator | 0.2 |
| $r_{p}$ | Reproductive cost on the predator due to parasite infection | [0-1] |
| $r_{e}$ | Reproductive cost on the predator due to parasite exposure | 1 |
| $d_{y}$ | The intrinsic death rate of the predator | 1 |
| $\mu_{y}$ | The mutation rate of the predator (per locus per reproduction) | 5x10^-5^ |
| $\sigma$ | Relationship between probabilities of successful infection and host resistance | 0.85 |

**Supplementary Note 2: Probability of successful infection**

In the following section, we explain how we calculate the probability of successful infection between a parasite and a prey $Q_{x}$. However, the same principle applies to the probability of successful infection between a parasite and a predator $Q_{y}=1$.

If the prey genotype $X^{j}$ at the $p$-$th$ loci has a resistant allele, we denote it as ${B_{p}(X}^{j})=1$. Similarly, ${B_{p}(Z}^{k})$ = 0 means the parasite genotype $Z^{k}$ at the $p$-$th$ loci has a non-infective allele. The effective resistance of the prey genotype *j* to parasite genotype $k$ is the sum of the matching resistant/non-infective alleles.

$r\left( X^{j},Z^{k} \right)= \sum_{p=1}^{n} {B_{p}(X}^{j})(1-{B_{p}(Z}^{k}))$	(1),

The probability of successful infection $Q_{x}$ will be determined by the effective resistance $r$ as:

${Q_{x}(X^{j},Z^{k})= \sigma}^{r(X^{j},Z^{k})}$	(2).

According to eq. 1, the maximum value $r$ is the number of loci $n$ (i.e., the length of the genotype), and the minimum value $r$ is 0, in a single host/parasite interaction. The concrete value of $r$ changes according to the number of effective resistant alleles (i.e., the number of resistant alleles matching non-infective alleles), which is also linked to the length of the genotypes $n$ (Figure S3). $\sigma$ is a parameter we input to the system, where different $\sigma$ values lead to a different relationship between the probability of successful infection $Q_{x}$(or $Q_{y}$) with $r$ (Figure S4). If $0<$ $\sigma<1$ then $Q_{x}$ (or $Q_{y}$) decreases with increasing $r.$ If $\sigma>1$, then $Q_{x}$ (or $Q_{y}$) increases with increasing $r$, which conflicts with our logic that increasing the number of effective resistant alleles leads to a lower probability of successful infection. Thus, we only use the parameter range$0<$ $\sigma<1$, which automatically guarantees the probability of successful infection, $0<$ $Q_{x}$ $<$ 1. The value of σ is assigned arbitrarily and can be changed to look at different outcomes from the simulations. We chose $\sigma=0.85$ to ensure that values of $Q_{x}$ and $Q_{y}$ changed gradually accordingly to the effective resistance of the prey and predator hosts, respectively (Figure S4).

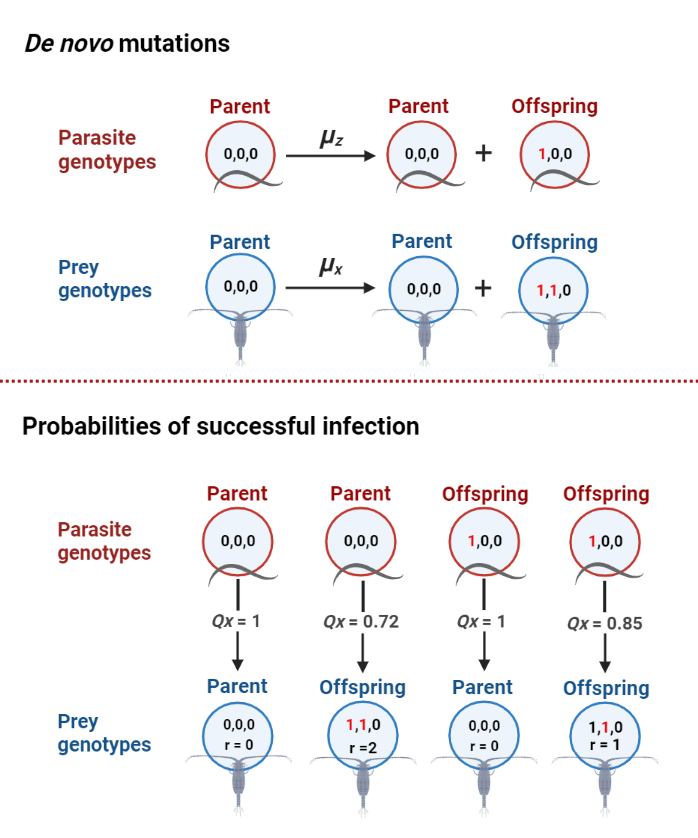

Figure S3. Probabilities of successful infection according to genotypic changes with *de novo* mutations. When a reproduction event happens, the genotype of the offspring may carry a different genotype than its parent, according to a mutation rate (top side of the figure). The probabilities of successful infection change according to the allelic states of the prey and parasite genotypes (bottom side of the figure). *r* is the sum of resistant alleles in the prey genotype (‘1’) matching the same position as non-infective alleles (‘0’) in the parasite genotype. The effective resistance *r* determines the probability of successful infection $\boldsymbol{Q}_{\boldsymbol{x}}$ according to ${\boldsymbol{Q}_{\boldsymbol{x}}\boldsymbol{= \sigma}}^{\boldsymbol{r}}$ where $\boldsymbol{\sigma=0.85}$. This illustration shows the probabilities of successful infection between a parasite and a prey. However, the same principle applies to parasites and predators. Note that this example illustrates the interaction of host and parasite genotypes of 3 loci. However, in our system, host (predator and prey) and parasite genotypes have 10 loci.

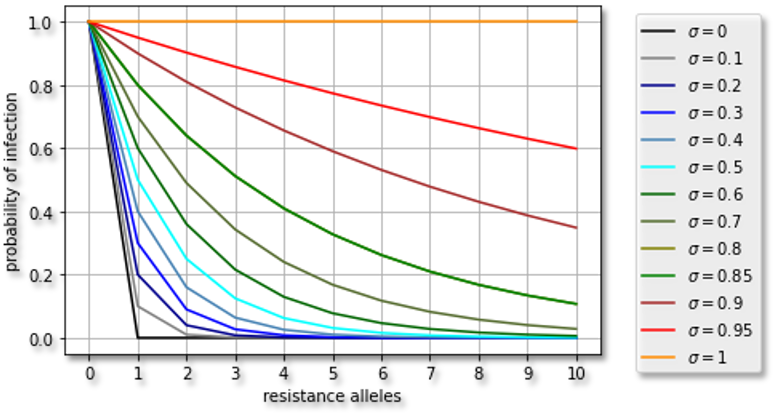

**Figure S4. Exponential distribution of probabilities of successful infection according to increasing values of σ.** The distributions of the probabilities of successful infection are described for host and parasite genotypes of 10 loci, as modelled in our system.

**Supplementary Note 3: Choice of time points for calculating the mean population size and genotypic diversity of each entity**

The correlations between predator, prey and parasite populations are different in punctual time points. The average of the 100 realisations is stable starting from time point 200-1000, and the distribution of different realisations fluctuates over time (Figure S5). This is consistent across different parameter sets; therefore, we used the average of 100 realisations from time point 200-1000 to represent the population size and genotypic diversity in each parameter combination of $r_{x}$ and $r_{p}$.

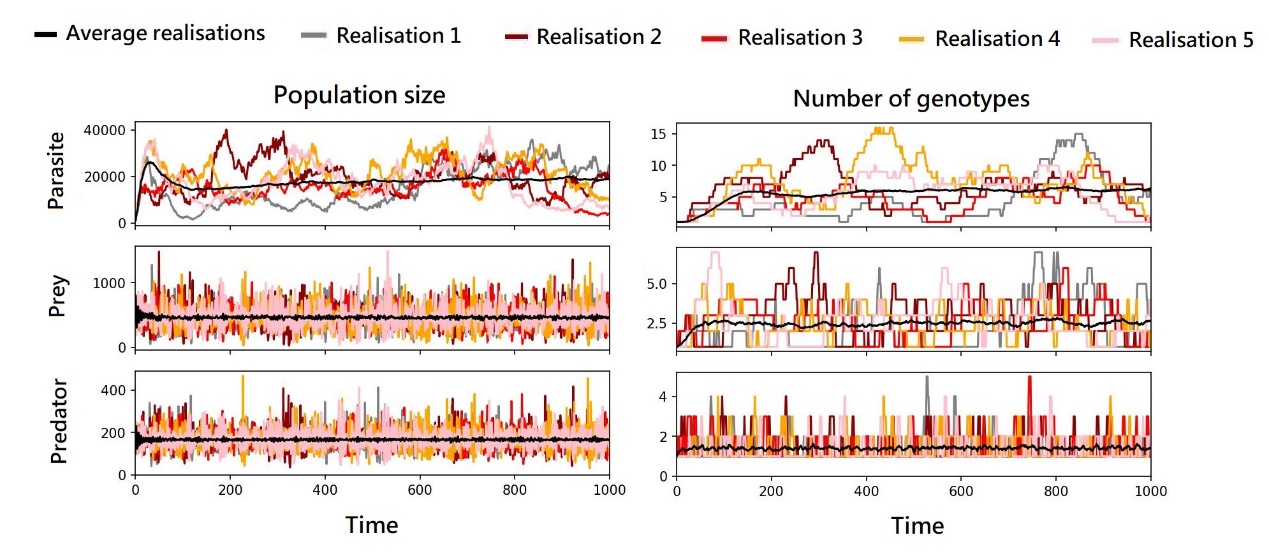

**Figure S5. Mean population size and genotypic diversity from the average of 100 independent realisations.** Parameter space in the absence of reproductive costs on the predator and prey. The top panels show the population size and the number of genotypes of the parasite, the middle panels show the population size and the number of genotypes of the prey, and the bottom panels show the population size and the number of genotypes of the predator. The left side panels show the population size and the right side panels show the number of genotypes in each time point for each entity. The black line shows the average from 100 independent realisations, and the color lines show the independent realisations fluctuation around the mean.

**Supplementary Figures**

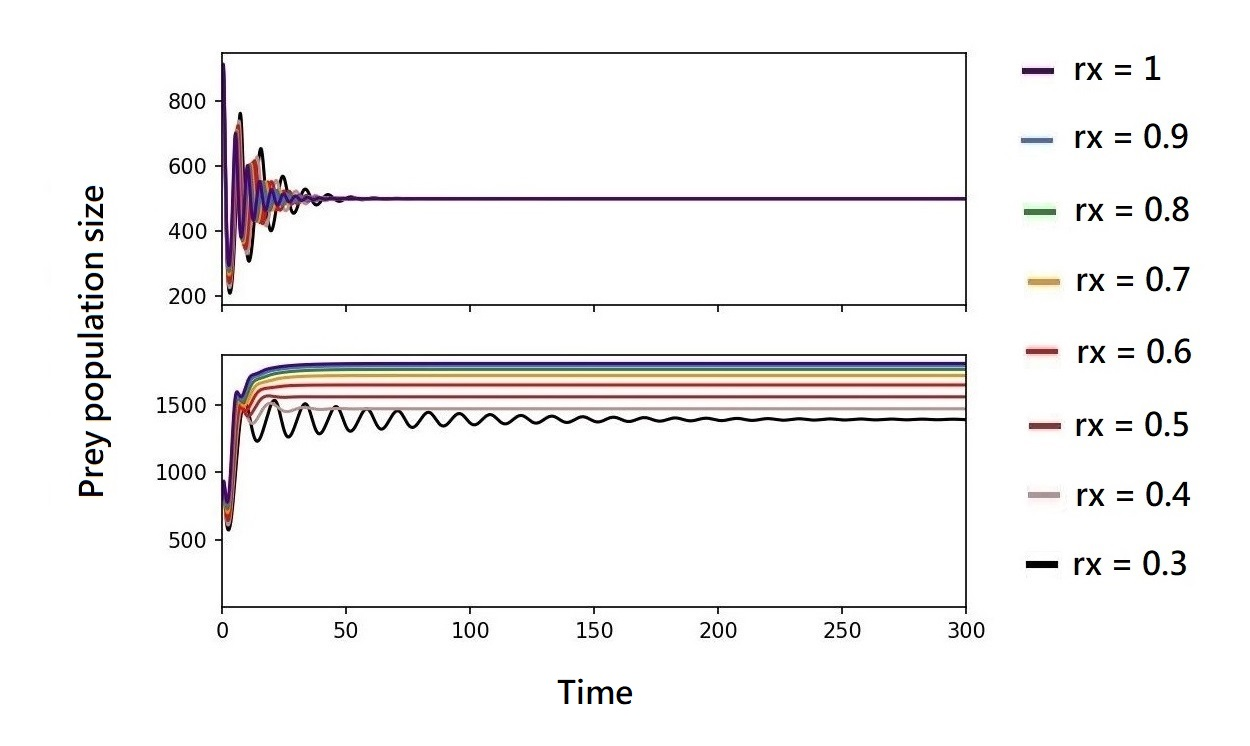
 **Figure S6. Equilibrium points of prey population dynamics under different selection pressures on the prey and predator.** The colors show the prey population dynamics according to different reproductive costs on the prey (i.e., $r_{x}$ values). The top panel shows prey population dynamics according to no parasite-mediated selection on the predator, $r_{p}$ = 1, i.e., predators are abundant; the bottom panel shows prey population dynamics according to intermediate parasite-mediated selection on the predator, $r_{p}$ = 0.5, i.e., predators are less abundant. Note that the values of $r_{x}$ and $r_{p}$ are opposite to the level of reproductive costs, where large values of $r_{x}$ and $r_{p}$ represent low reproductive costs.

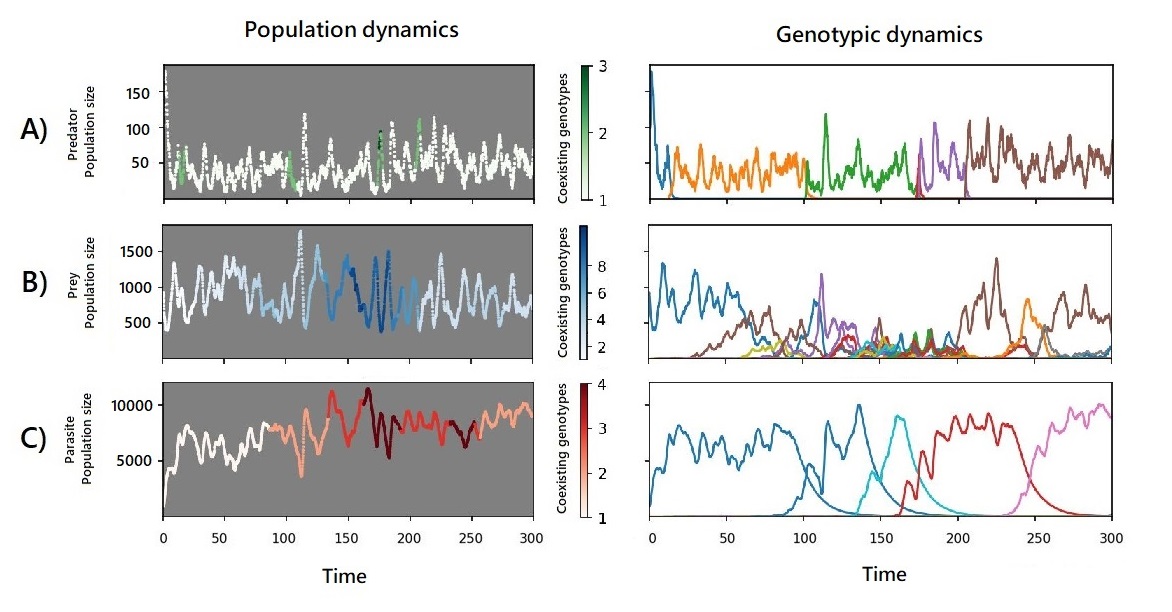

**Figure S7.** **Population and genotypic dynamics under high reproductive costs on the prey and low reproductive costs on the predator, from a single realisation.** The left panels show the population dynamics of the species, where darker shading indicates a higher number of genotypes coexisting. The right panels show the genotypic dynamics of the species, where different colors show the emergence of different genotypes. A) population and genotypic dynamics of the predator, B) population and genotypic dynamics of the prey, and C) population and genotypic dynamics of the parasite. Parameter values: $r_{x}=0.3$ and $r_{p}=0.7$.

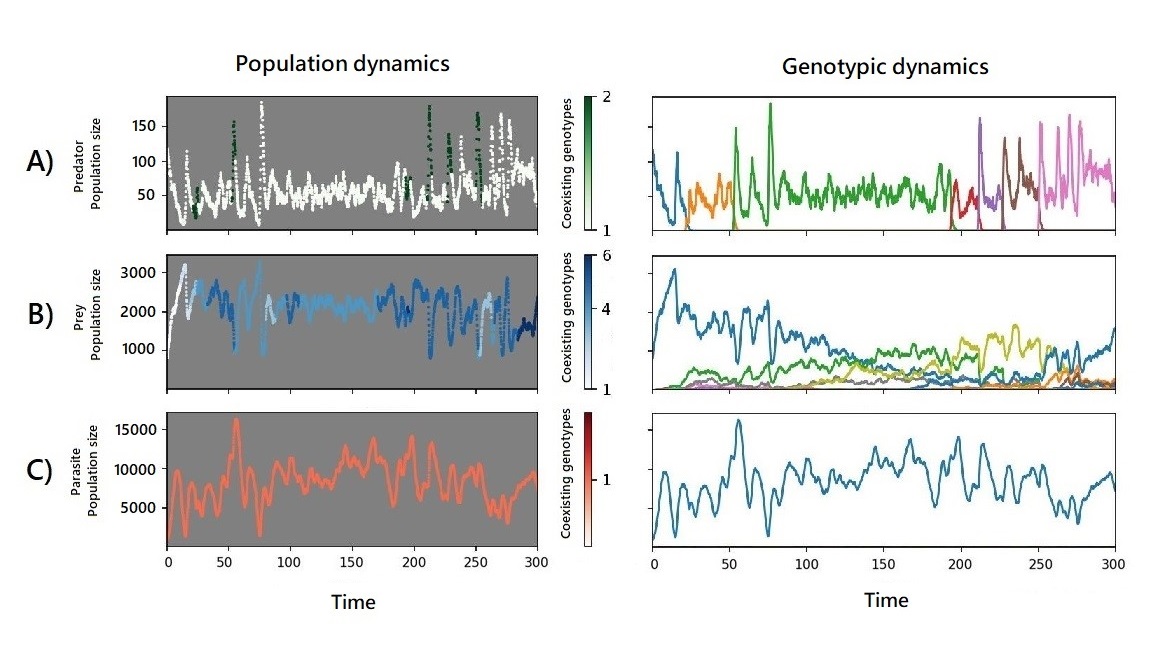

**Figure S8.** **Population and genotypic dynamics under low reproductive costs on the prey and high reproductive costs on the predator, from a single realisation.** The left panels show the population dynamics of the species, where darker shading indicates a higher number of genotypes coexisting. The right panels show the genotypic dynamics of the species, where different colors show the emergence of different genotypes. A) population and genotypic dynamics of the predator, B) population and genotypic dynamics of the prey, and C) population and genotypic dynamics of the parasite. Parameter values: $r_{x}=0.8$ and $r_{p}=0.3$.

**Supplementary Tables**

**Table S2.** **Correlations between mean population size of predators, prey, and parasites as a function of parasite-mediated selection on the prey and the predator.** A) Correlation between parasite and prey mean population size B) Correlation between predator and parasite mean population size C) Correlation between predator and prey mean population size. The panels show different degrees of parasite-mediated selection on the predator $r_{p}$, where smaller values indicate higher selection. The colored dots indicate the magnitude of parasite-mediated selection of the prey $r_{x}$, where smaller values indicate higher selection. Each colored point displays the average value from 100 independent realisations over a time period of 200-1000 time steps for each realisation.

| **A)** 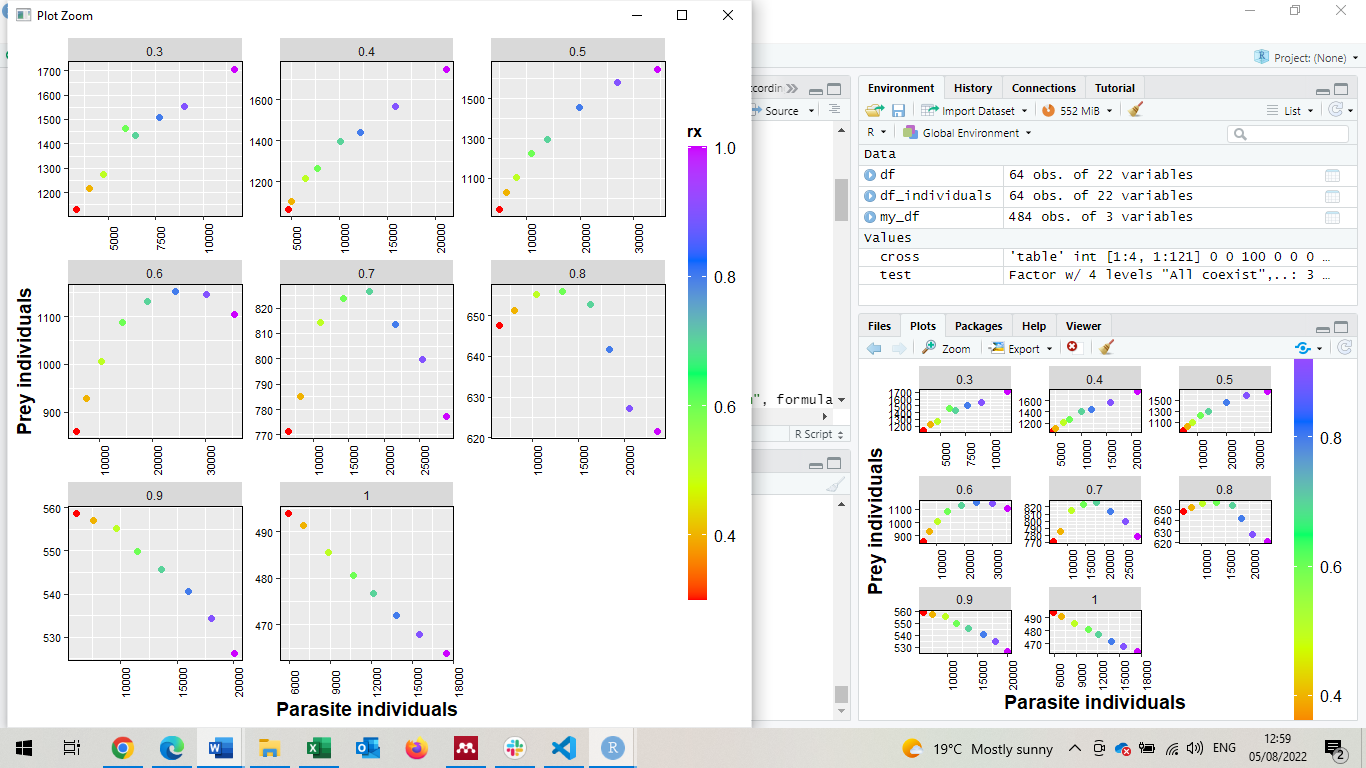 | $r_{p}$ = 0.3: F_2,5_ = 76.95; p = 0.0001  $r_{p}$ = 0.4: F_2,5_ = 274.7; p < 0.0001  $r_{p}$ = 0.5: F_2,5_ = 929.6; p < 0.0001  $r_{p}$ = 0.6: F_2,5_ = 258.5; p < 0.0001  $r_{p}$ = 0.7: F_2,5_ = 67.45; p = 0.0002  $r_{p}$ = 0.8: F_2,5_ = 37.06; p = 0.001  $r_{p}$ = 0.9: F_2,5_ = 1,117; p < 0.0001  $r_{p}$ = 1: F_2,5_ = 8,778; p < 0.0001 |
| --- | --- |
| **B)** 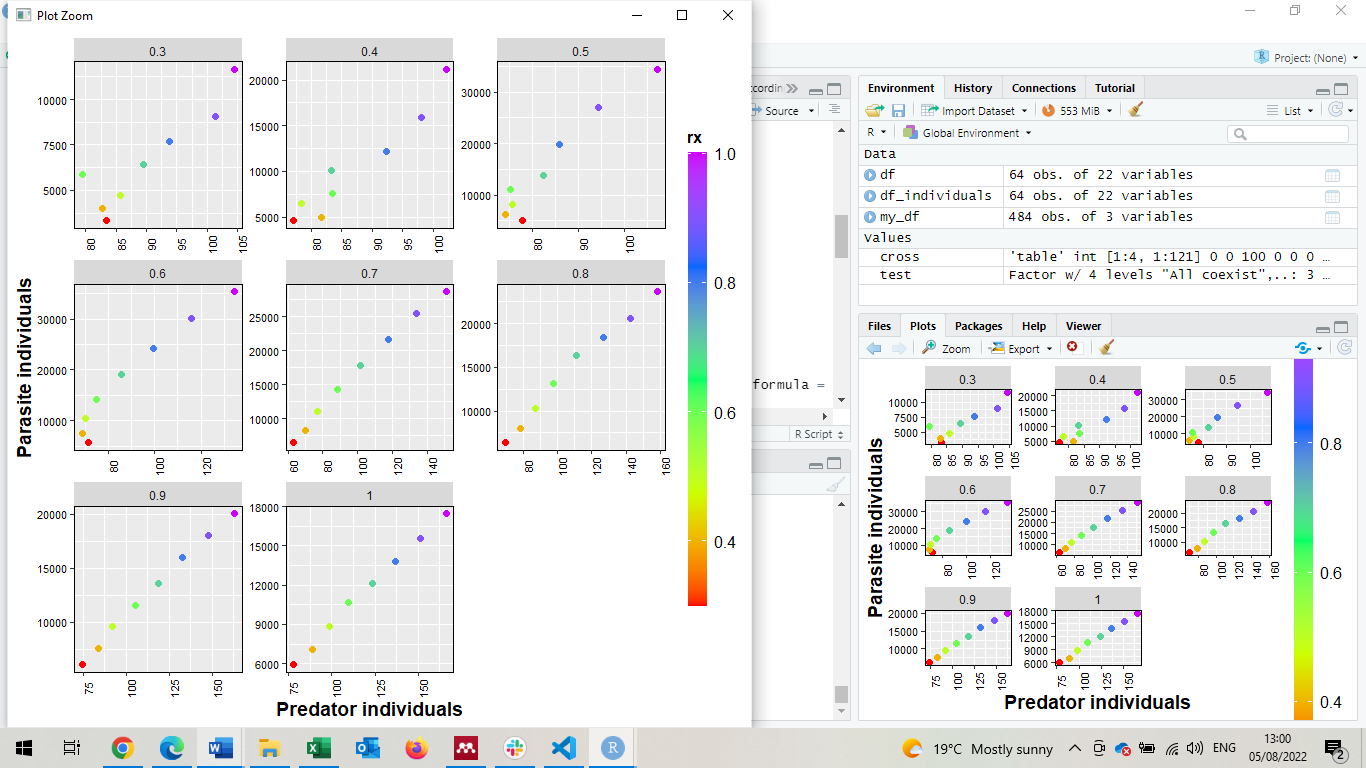 | $r_{p}$ = 0.3: F_2,5_ = 18.93; p = 0.0046  $r_{p}$ = 0.4: F_2,5_ = 45.03; p = 0.0006  $r_{p}$ = 0.5: F_2,5_ = 41.33; p = 0.0007  $r_{p}$ = 0.6: F_2,5_ = 74.24; p = 0.0002  $r_{p}$ = 0.7: F_2,5_ = 4,442; p < 0.0001  $r_{p}$ = 0.8: F_2,5_ = 493.2; p < 0.0001  $r_{p}$ = 0.9: F_2,5_ = 2,005; p < 0.0001  $r_{p}$ = 1: F_2,5_ = 1,095; p < 0.0001 |
| **C)** 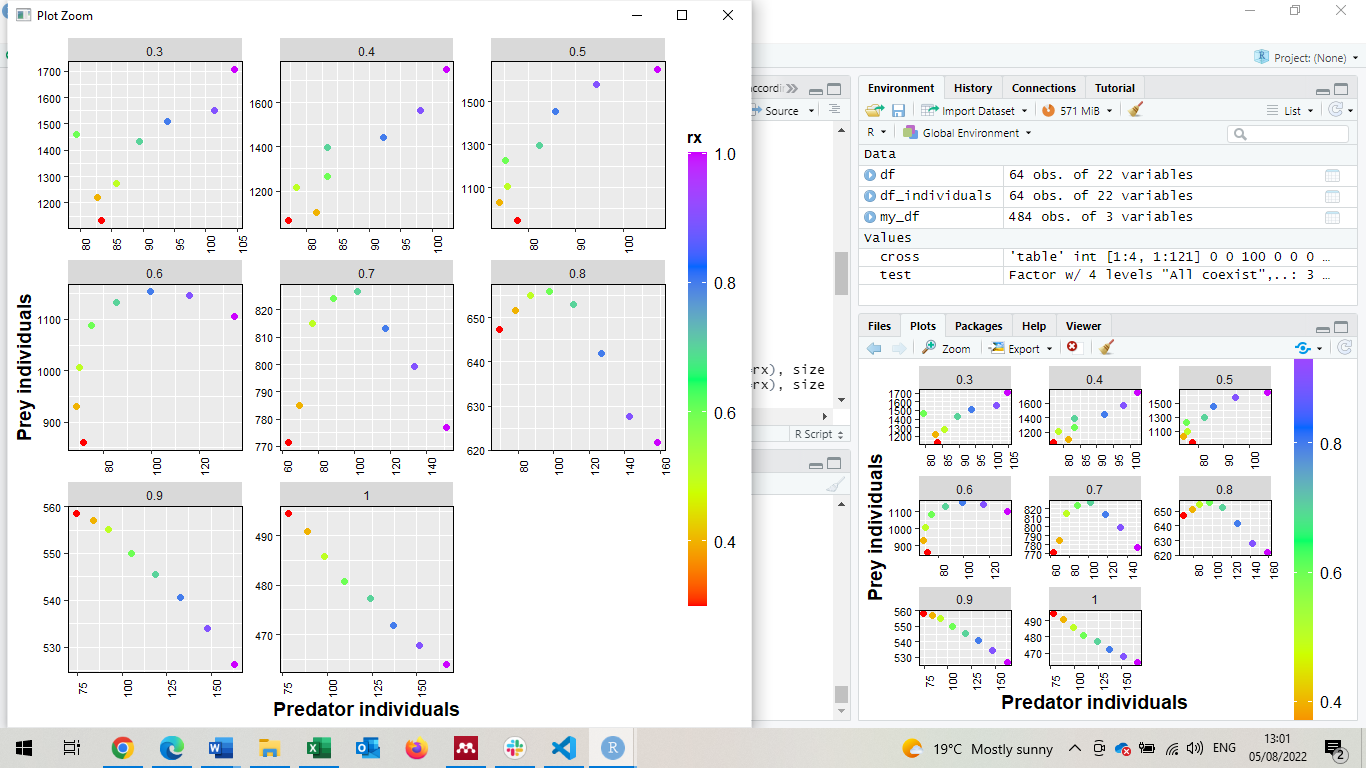 | $r_{p}$ = 0.3: F_2,5_ = 5.351; p = 0.0572  $r_{p}$ = 0.4: F_2,5_ = 19.16; p = 0.0045  $r_{p}$ = 0.5: F_2,5_ = 13.25; p = 0.01  $r_{p}$ = 0.6: F_2,5_ = 6.139; p = 0.045  $r_{p}$ = 0.7: F_2,5_ = 18.59; p = 0.0048  $r_{p}$ = 0.8: F_2,5_ = 45.18; p = 0.0006  $r_{p}$ = 0.9: F_2,5_ = 3,815; p < 0.0001  $r_{p}$ = 1: F_2,5_ = 984.6; p < 0.0001 |
